# Artisanal and farmer bread making practices differently shape fungal species community composition in French sourdoughs

**DOI:** 10.1101/679472

**Authors:** Elisa Michel, Estelle Masson, Sandrine Bubbendorf, Léocadie Lapicque, Thibault Nidelet, Diego Segond, Stéphane Guézenec, Thérèse Marlin, Hugo Devillers, Olivier Rué, Bernard Onno, Judith Legrand, Delphine Sicard, the participating bakers

## Abstract

Preserving microbial diversity in food systems is one of the many challenges to be met to achieve food security and quality. Although industrialization led to the selection and spread of specific fermenting microbial strains, there are still ongoing artisanal processes that may allow the conservation of a wider species diversity and genetic diversity. We examined whether the diversity of artisanal practices could lead to an increased level in fungal species diversity for bread making. We used an interdisciplinary participatory research approach including bakers, psycho-sociologists and microbiologists to analyze French bread making practices and describe fungal communities in naturally fermented sourdough of 27 bakers and 12 farmer bakers. Bread making practices were classified in two groups: the farmer-like practice group and the artisanal-like practice group. The well-known bakery yeast, *Saccharomyces cerevisiae*, was dominant (i.e. with a relative abundance over 50%) in only 24% of sourdoughs while other yeast species, belonging to the *Kazachstania* genus, were dominant in 54% of sourdoughs. Bread making practices were found to drive the distribution of fungal species across sourdoughs. The most striking bread making practice effect was the occurrence of *Kazachstania humilis* in sourdoughs made with artisanal-like practices and the occurrence of *Kazachstania bulderi* in sourdoughs made with farmer-like practices. Phenotypic divergences between sourdough and non-sourdough strains were found for *K. humilis* but not for *K. bulderi*. Overall, our results showed that preserving bread making practice diversity allows the preservation of a higher species and phenotypic diversity in microbial communities.

## Introduction

Humans started to ferment food before the Neolithic using naturally fermenting microbial communities. In the 19^th^ century, the industrialization and the increase of knowledge in microbiology resulted in changes in fermented food practices with the use of starters. This selection led to a reduction in species diversity and genetic diversity for fermented food processing and limited *in situ* conservation of microbial communities in industrialized systems [1–3]. Domestication of the yeast *Saccharomyces cerevisiae* for the production of beer, wine, cheese, leavened bread, that of the fungi *Penicillium roqueforti* or *Penicillium camemberti* for cheese production or that of the fungus *Aspergillus oryzae* for rice or soybean fermented products are well studied cases [1–10]. The recent renewed interest in artisanal practices that make use of naturally fermenting microbial communities could promote the conservation of microbial diversity. However, the effect of artisanal practices on the distribution of microbial species across sourdoughs remains poorly documented.

Among fermented foods, bread is still a symbol deeply engrained in the history, religious rites and medicine of several cultures. Bread likely originated 14 000 years ago, suggesting that bread was made long before plant domestication [11]. Since the Neolithic, bread history is intimately associated with the domestication of cereals, bread making associated tools and the advent of Mediterranean civilizations [12]. Investigation of the morphology of plant remains which were incorporated in Neolithic bread identified wheat, barley, millet, linseed [12]. Leavened bread was traditionally made with flour, water and a fermenting agent, which was either a fermenting beverage or a fermenting dough, called sourdough. This sourdough was generally initiated from a mixture of flour and water, naturally colonized by lactic acid bacteria (LAB) and yeasts. Sourdough was then either maintained over time or initiated again and again, depending on the craftsman [13, 14]. In the 19^th^ century, the use of yeast starters made of *S. cerevisiae*, often called « baker’s yeast », spread as an alternative to sourdough. Nowadays, *S. cerevisiae* industrial starters are more frequently used than sourdoughs, although the latter are gaining interest. A recent study showed that industrial populations of *S. cerevisiae* have followed a different evolutionary path than sourdough populations [10]. Both have been domesticated by humans which has improved their fermentation performance in a sourdough-mimicking medium. Industrial and sourdough strains of *S. cerevisiae* differ genetically and phenotypically, indicating that sourdough use contributes to the conservation of bread related *S. cerevisiae* lineages [10].

Yeasts, are organisms growing mainly as single cells and with sexual states not enclosed in fruiting bodies, belonging to ascomycete or basidiomycete fungi. To date, more than 40 yeast species have been detected in sourdough [1, 15, 16]. The most frequently encountered species are *Wickerhamomyces anomalus* and *Kazachstania humilis*. Several other species in the genus *Kazachstania (Kazachstania barnettii, Kazachatania exigua, Kazachstania bulderi, Kazachstania unispora*) as well as several species in the polyphyletic genus *Pichia* have also been recurrently detected. The factors determining the presence in sourdough of these species are still unknown. A recent large-scale study of 500 sourdoughs from four continents found no effect of geography or factors related to bread making practices such as age of sourdough, storage location, feeding frequency, or grain intake [17]. However, most of the sourdoughs in this study were made by private citizens who probably did not maintain the sourdough microbial community in the same way as professional bakers. To our knowledge, no studies have been conducted to date to investigate the effect of bakers’ bread making practices on sourdough yeast community composition.

In France, sourdough breads are made both by bakers and farmers who also grow and mill their own wheat. The number of farmer-bakers has increased in the 2000s with two motives: to grow wheat varieties meeting their needs and to assert their independence from industry [18]. Although farmer-bakers are less numerous than bakers, they participate in the renewed interest in local wheat varieties and artisanal knowhow, which may contribute to the conservation of both socio-cultural diversity and microbial diversity.

Here, we used a participatory research approach involving psycho-sociologists, biologists, biostatisticians, bakers and farmer-bakers to study whether and how bakers and farmer-bakers contribute to the preservation of socio-cultural and fungal species diversity in sourdough microbial community.

## Methods

A total of 27 bakers and 12 farmer-bakers participated to the study. They were all making bread with organic flour except five. All of them sent sourdough to the lab for microbiological and metabolic analysis. Among them, 36 described their bread making practices as well.

### A questionnaire survey, face-to-face interviews and focus-groups to collect bread making practices

Data on bread making practices were collected through a questionnaire survey, interviews and focus groups. The collected variables were related to *i*) the ingredients origin: wheat varieties types (ancient populations also called landraces / modern varieties), whether they produced flour from their own wheat, whether they had their own mill or use an external mill, water origin, *ii*) the sourdough recipe: its age, its hydration state, the origin of the chief sourdough (sample of dough or sourdough), the number of back-sloppings before bread making and per week, the temperature of water used for back-sloppings, *iii*) their bread making practices: the number of bread makings per week, the percentage of sourdough, flour and salt in bread dough, the kneading methods, the total duration of fermentation and the addition of baker’s yeast in dough.

### Sourdough samples, enumeration and strain isolation

Sourdoughs were collected before kneading and referred to as final sourdoughs (Table S1 [87]). On the day of collection, they were sent to the lab where yeasts and bacteria were enumerated and isolated as in [19, 20], and sourdoughs stored at −20°C in sterile vials for non-culture based analysis. Ethics and rights associated with sourdough collection and strain isolation have been respected.

### Sourdough acidity and metabolic analyses

For each sourdough, three independent 1-g replicates were analyzed. pH and Total Titrable Acidity were measured as described in [20]. Organic acids, alcohol and sugar concentrations (expressed as g/kg of sourdough) were analyzed by liquid chromatography using an HPLC HP 1100 LC system (Agilent technologies, Santa clara, CA, USA) equipped with a refractive index detector (RID Agilent G1382A) and a UV detector (Agilent G1314A). Two different columns were used, a Rezex ROA-organic acid column and a Rezex RPM-monosaccharide column (SDVB – Pb+2 8%, 300×7.8mm, Phenomenex, Torrance, CA, USA). The details of the experiments are described in supplementary information (Method S1 [87]).

### Yeast species identification

The Internal transcribed spacer 1 (ITS1) ribosomal DNA of each 1216 yeast isolates was amplified by PCR from chromosomal DNA, either by using primers ITS1F and ITS2 [21, 22], or primers NSA3 and 58A2R [19, 22]. For isolates unidentified with the ITS1 region alone, DNA was extracted with the MasterPure yeast DNA purification kit (Epicentre, Epibio). PCR reactions targeting partial genes, the D1D2 region of the large subunit of rRNA (LSU), a part of the RNA polymerase II large subunit encoding gene (*RPB1*), a part of the RNA polymerase II encoding gene (*RPB2*), a part of the actin encoding gene (*ACT1*) and transcriptional elongation factor (*TEF alpha*) were performed. To discriminate three specific isolates, PCR on genes *GHD1, FSY1, URA3, DRC1, MET2* were performed [23–26] (Table S2 [87]). All PCR products were sent to be sequenced with Sanger sequencing (Eurofins, Germany). Species were identified using NCBI [27], YeastIP [28] and a personal database, which was constructed after *ITS1, RPB2*, LSU sequencing of all 33 yeast species reportedly found in sourdoughs in the literature [19].

### Sourdough DNA extraction, MiSeq sequencing, bioinformatics

The ITS1 region was targeted with the PCR primers ITS1F (5’-CTTGGTCATTTAGAGGAAGTAA - 3’) and ITS2 (5’-GCTGCGTTCTTCATCGATGC-3’).

The sequencing run was performed with MiSeq Reagent Kit v3. 2015 [20]. Sequences were analyzed through FROGS “Find Rapidly OTU with Galaxy Solution” [29] and home-made pipelines. Overlapped reads were merged with Flash [30] with a minimum overlap of 10 nucleotides, a maximum overlap of 300 nucleotides and a maximum mismatch density of 0.1. Primers were removed with Cutadapt [31] and data were cleaned with Sickle with quality-threshold and length-threshold equal to 20 [32]. Reads were clustered with Swarm (d=3) [33] and chimeras deleted with VSEARCH [34]. Sequences were then filtered on minimum abundance of 0.005% of all sequences. From the OTU abundance table and for each OTU, the taxonomic affiliation using UNITE Version 7.1, Release 2016-11-20 [35], YeastIP [28] and our own database [19] was obtained by blasting operational taxonomic units (OTUs) representative sequences against each database.

### Phenotypic analysis of yeast strains

Fermentation performance of the two most frequently encountered *Kazachstania* species was assessed as described in [10] for *S. cerevisiae*. Fifteen sourdough strains of *K. bulderi* and 16 sourdough strains of *K. humilis* were included in the analysis. *Kazachstania bulderi* strains were coming from sourdoughs B4, B12, B15, B17, B20, B21, while *K. humilis* strains were coming from sourdoughs B2, B5, B6, B7, B10, B17. From one to three strains per sourdough were analyzed in the experiment. In addition, four strains of *K. bulderi* (strain MUCL 38021 isolated from silage in Namur, Belgium, strain MUCL 54694 isolated from silage in Erezée, Belgium, strain NRRL Y-27205 and strain CLIB 604 isolated from maize silage in the Netherland) and three strains of *K. humilis* (strain CBS 7754 isolated from food dressing in Germany, strain CLIB 1323 isolated from bantu beer in South Africa, and strain CBS 2664 isolated from alpechin in Spain) which were coming from non-sourdough habitats, were added as control to test the effect of habitat of origin. Each strain was phenotyped at least in triplicate leading to a total of 145 fermentations distributed over two blocks. Briefly, fermentations were carried out at 24 °C with constant magnetic stirring (300 rpm) during 24 h. CO_2_ release was measured by weight loss every 40 min using an automated robotic system [36]. At the end of fermentation, population size and cell viability were determined by flow cytometer (C6 cytometer, Accuri, BD Biosciences) as described in [37].

### Data analyses

A multiple correspondence analysis (MCA) and hierarchical clustering (complete linkage clustering method) on principal components based on the first two axes of the MCA were performed using the FactoMineR R package [38].

To analyse fungal community, weighted UniFrac distances between sourdough communities were computed from a rooted phylogenetic tree based on the OTU sequences using the R-packages Phyloseq and GUniFrac [39, 40]. Phylogenetic sequences were aligned with Clustal Omega and phylogenetic trees were built with the parsimony algorithm, with 100 replicate bootstraps, pairwise ktuple-distances with Seaview [41]. The results presented in the main text were obtained using the phylogenetic tree rooted on the OTU identified as *Sporidiobolales* species. Different roots were tested. The roots were chosen among the OTUs that were affiliated to the most distant taxa (*Sporidiobolales sp., Bullera globospora, Trichosporon asahii, Udeniomyces pyricola*). The tree topology did not change with the chosen root. The tree did not fit the expected phylogeny and, notably, some *Ascomycota* were located among the *Basidiomycota*. However, the dominant sourdough species belonging to the *Saccharomycetaceae* family were clustered in the expected clades or subclades, except that *Kazachstania servazzii* and *Kazachstania unispora* were grouped in a clade closer to *Saccharomyces* species than to other *Kazachstania* species. Using the UniFrac distances matrix, we performed a principal coordinate analysis (PCoA) and clustered sourdough communities using the first two axes of the PCoA, and the complete linkage clustering method (hclust R function). To check the sensitivity of our analysis to this misclassification, we performed the same analyses without the sourdoughs that had one misclassified species representing more than 10% of their reads, *i.e*. sourdoughs B20, B41, B42, and B44 and found the same clustering [40].

For each sourdough, the species richness, Chao1, Shannon and Simpson indexes were computed. Chao1 was used as an indicator of species richness corrected by the number of OTUs present in the community but not observed. Shannon and Simpson index values were converted to the effective number of species per sourdough. This number was estimated from the Shannon diversity index as *exp^shannonindex^* and from the Simpson diversity index as 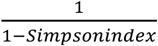 [42, 43]. For probability estimates, the exact 95% confidence intervals were computed using a binomial distribution.

To investigate the relationship between β-diversity and differences in bakery practices, we performed a permutational multivariate analysis of variance (PERMANOVA) on the UniFrac distance matrix for each bakery practice variable. We included in the analysis sourdough fungal communities of the 30 bakers who had less than 8 missing values among the 29 bread making practices variables and adjusted the p-values using false discovery rate method correction to account for multiple testing [44]. In addition, we performed independence exact Fisher tests between fungal community PCoA groups and each of the bread making practices variables. Multiple testing was accounted for using the false discovery rate method [44].

In addition, we tested the link between the baker practice group, the fungal community group or the yeast dominant species and the variation of each quantitative variable (microbial density, pH, TTA, metabolite concentration) with the following mixed effect model: *Y_ijk_* = μ + *α_i_* + *B_j_* + *ε_ijk_* with *ε_ijk_* ~N(0, σ^2^), where *α_i_* is the effect of the fungal community group *i* modelled as a fixed effect and *B_j_* is the effect of sourdough j modelled as a random effect and k represents the measurement replicates. For sourdough hydration rate, the variable was arcsin transformed but sourdough effect was not included in the model because no repetition was obtained from any sourdough. The model parameters were estimated using the lmertest R package [45]. To test the fixed effects, we used likelihood ratio tests. Multiple comparisons of means were performed using Tukey tests with the multcomp package. p-values were all adjusted for multiple testing with the FDR method. The geographical structuration was tested with a Mantel test on the UniFrac distances matrix and the geographical distances matrix computed with the package geosphere [46] and ade4 [47].

The phenotypic diversity of *K. bulderi* and *K. humilis* strains coming from sourdough and non-sourdough habitats were analyzed. Population size and mortality rate after 27h of fermentation measured by flow cytometer were used as proxies for absolute fitness. The cumulative CO_2_ production curve was calculated and the kinetics of CO_2_ production rate over time was estimated by successive linear smoothing over five points. Four fermentation parameters were then estimated. The maximum CO_2_ release (CO_2_max, in g/L) was estimated by the maximum of the cumulative CO_2_ production curve. The fermentation latency phase time was estimated by the time between inoculation and the beginning of the fermentation calculated as 1g/L of CO_2_ release (t1g, in h). The maximum CO_2_ production rate (Vmax in g/L/h) was estimated by the maximum of the CO_2_ production rate kinetic. The time of the Vmax (tVmax in h) was calculated as the time between inoculation and the Vmax. Hence, the phenotype of each strain was characterized by six quantitative variables called “phenotype variables” below: its population size, its mortality rate and the four fermentation parameters. To determine whether the origin of the strain (sourdough or non-sourdough) had an impact on strain phenotype, each log-transformed quantitative variable was analyzed separately using a mixed linear model as described below. The experimental design was unbalanced between the two blocks with very few non-sourdough strains in one of the two blocks. Therefore, for each phenotype variable, we first estimated the block effect with a subset of 8 strains cultivated in both blocks using a linear model with two fixed effects: the strain and the block. Additive models were used as the interaction terms were not significantly different from zero after adjusting p-values with the Benjamini-Hochberg method. Second, each phenotype variable was corrected for the block effect and analyzed with the mixed effect model: *Y_ijkl_* = μ + *α_i_* + *β_j_* + *γ_ij_* + *Z_k_* + *ε_ijkl_* where *Y_ijkl_* represents the log-transformed phenotype variable corrected for the block effect for the strain *k*, from species *i (i=1,2)*, sampled in environment *j (j=1,2)*, observed for replicate *l. μ* represents the mean of the phenotype variable, *α_i_* the additive effect of species *i, β_j_* the additive effect of environment *j*, and *γ_ij_* their interaction. *Z_k_* represents the gaussian random effect of strain *k* with 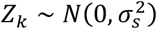 and *ε_ijkl_* the gaussian residuals with *ε_ijkl_* ~*N*(0, σ^2^). For each species *i*, the impact of the environment was quantified using the contrast *Δ_i_* = *β_s_* + *γ_i,s_* – *β_KS_* – *γ_i,NS_* with “NS” standing for “non-sourdough” and “S” for “sourdough” and tests (*H*_0_ = Δ*_i_* = O, *H*_1_:*Δ_i_* ≠ 0) were performed and p-values were adjusted using the Benjamini-Hochberg method. As log-transformed data were analyzed, the exponential of this contrast can be interpreted as the ratio between the sourdough mean and the non-sourdough mean. Confidence intervals and tests were performed using the doBy R package.

All statistics and plots have been done with R (ggplot2 [48], leaflet package [49], with minor esthetical adjustment with Inkscape). Data and scripts are shared on Zenodo: DOI: 10.5281/zenodo.5849058 [87].

## Results

### Two groups of bread making practices

A total of 39 French bakers producing natural sourdough bread and distributed all over France participated to the study (Table S1 [87]). The bread making practices of 35 of them were collected through one or several methods: personal interviews (with 12 bakers), focus groups (three groups), observation during bread making workshops (two workshops), and an online/phone survey (35 bakers). The general process of sourdough bread making is presented in Figure 1. We analyzed 28 variables, describing variations of the practices at all steps of the bread making process, from wheat grains to baked bread (Figure S1 [87]). Four bakers (B6, B10, B19, B20) who did not provide enough information about their practices were excluded from the multivariate analysis.

**Figure 1:**
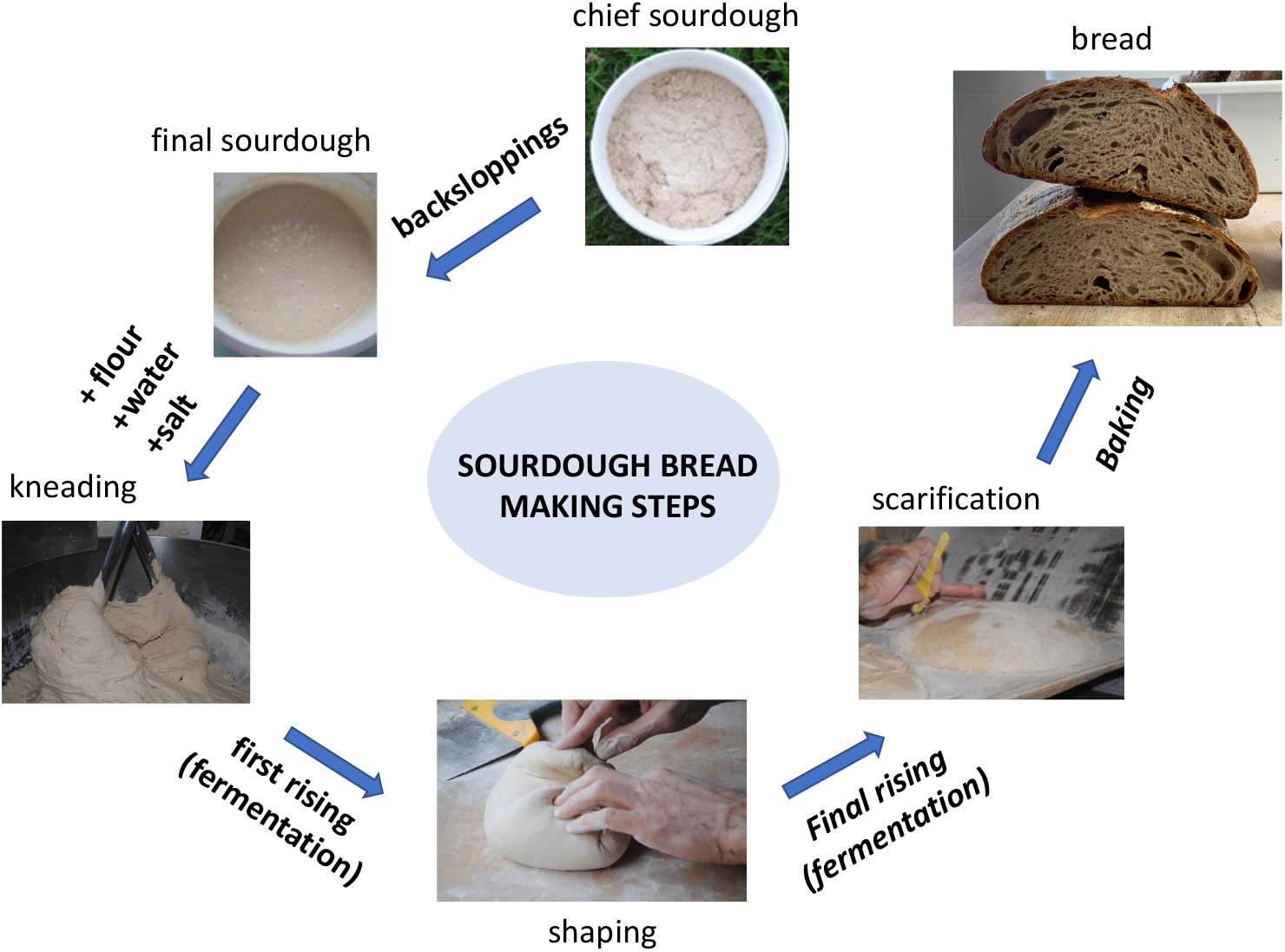
The sourdough bread making process. Sourdough is a mix of flour and water naturally fermented by bacteria and yeasts. It is initiated by mixing flour, water and occasionally other ingredients. It is then “fed” by regularly adding flour and water, a process termed back-slopping. Once considered mature by bakers based on their acidity, flavour and bubbling activity, the sourdough is called “chief”, or “mother” sourdough, and can then be used for bread making. The bread making process starts from this “chief sourdough”, or from a piece of dough or sourdough sampled from the preceding bread making process, or initiated from a mix of flour and water naturally colonized by yeasts and lactic acid bacteria following several back-sloppings. Once or several times, the chief sourdough is refreshed by adding flour and water to constitute the final sourdough, which is used for bread making. This final sourdough is mixed with flour, water, and other ingredients (salt, seeds, yeasts starters, etc.) during kneading to constitute the dough. After kneading, primary fermentation occurs during the first rising. The dough is then divided and shaped. The pieces of dough are then left to rise during a second fermentation and finally oven-baked.

According to a hierarchical clustering on principal components (HCPC), the 32 other bakers clustered into two groups corresponding to two main types of bread making practices (Figure 2). The first group, hereafter termed “farmer-like” practice group, included six bakers and 11 farmer-bakers using the following practices: low bread production (<500 kg per week, 81% of the bakers of the “farmer-like” group), use of wheat landraces (56%), manual kneading (63%), working at ambient temperature (88%), long fermentation periods (more than 4 hours for 88%), and no use of commercial baker’s yeast (88%). In addition, they tend to make their chief sourdough from dough after kneading (75%). The second group, hereafter called “artisanal-like” practice group consisted of 12 bakers and four farmer-bakers having more intensive practices, characterized by a large bread production (>500 kg per week, 81%), mechanical kneading (100%), use of modern wheat varieties (63%), working at ambient temperature (56%), using *S. cerevisiae* starters in addition to sourdough for bread making or for pastries and buns making (81%). In this second group, bakers tended to make their chief sourdough from a final sourdough.

**Figure 2:**
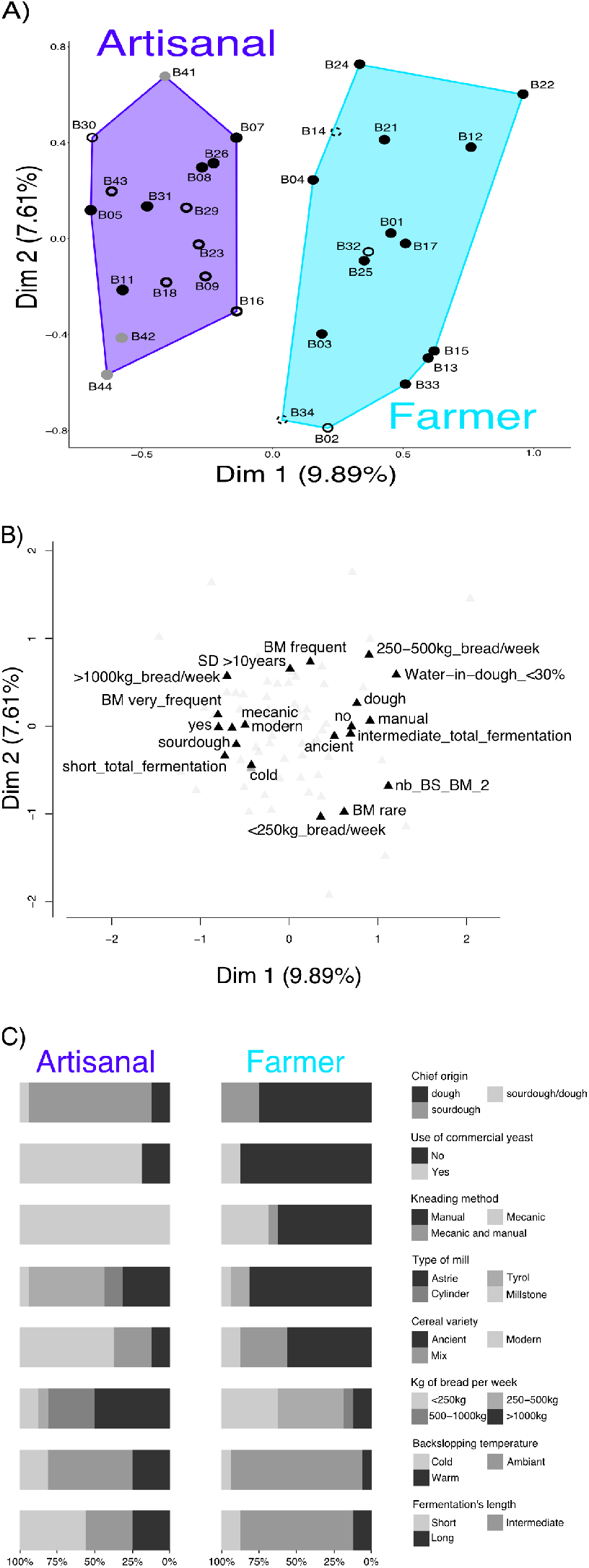
Multiple Correspondence Analysis (MCA) based on 28 categorical variables describing bread making practices. A) Representation of bakers. Each point represents a bakery. The purple area on the left brings together baker with “artisanal” practices and the light blue area on the right the bakers with “farmer” practices. The dot’s colors indicate the PCoA cluster of the sourdough fungal community (see Figure 5). Black dots for group 1, empty dot for group 2, grey for group 3. The fungal community of the sourdough of baker 14 was not studied. B) Representation of the 20 first categories that contributed the most to the MCA axes. The category, which corresponds to a class of a variable, is written next to the triangle. C) Distribution of each variable for each bread making practice group. Only variables that mostly explained differences between bread making practice groups are shown: use of commercial yeast, kneading method, chief origin, kg of bread production per week, number of bread making per week, percent of water in dough, number of back-sloppings before making bread, water origin, sourdough age and flour percentage in dough. The categories of each of these variables are indicated on the right.

### Composition of sourdough fungal communities

Sourdough is a mix of flour and water naturally fermented by bacteria and yeasts. Sourdough yeast density ranged from 8.1 10^4^ to 5.8 10^8^ CFU per gram of sourdough, with a mean value of 2.9 10^7^ CFU per gram, as commonly found in sourdoughs from all over the world [1, 16, 50, 51]. We isolated 20 to 40 yeast strains from each sourdough by picking colonies randomly and identified species using ITS sequence as well as other barcodes when the ITS alone was not able to discriminate between closely related species. Among the 39 collected sourdoughs, one (sourdough B14) did not give any colony in the laboratory, suggesting that his sourdough microbiota was no longer alive. A total of 1216 strains were characterized from the other 38 sourdoughs. In addition, we developed an ITS1 meta-barcoding MiSeq sequencing method on sourdough (see sup M&M). After filtering 5,360,620 raw ITS1 sequences for quality, abundance (0.005%) and chimera, 3,542,801 sequences were further analyzed. Overall, the sequences clustered in 113 OTUs. The number of reads per sourdough ranged from 8421 to 194,557. Therefore, we carried out our analysis on the rarefied matrix. Among all OTUs, 10 were assigned to the order *Triticodae* (especially to the species *Triticum aestivum*), 50 were assigned to a filamentous fungal genus including plant pathogen species such as *Alternaria, Aspergillus, Fusarium*, or *Gibberella*, while 4 OTUs remained unidentified. Among the 40 yeast OTUs, 96% of total reads were assigned to the phylum *Ascomycota*, 87.5% to the order *Saccharomycetales* and 85.7% to the family *Saccharomycetaceae*. Only 4% of the total reads were assigned to the phylum *Basidiomycota*. Overall, three OTUs assigned to the species *Kazachstania humilis, Kazachstania bulderi* and *Saccharomyces cerevisiae* represented 20.3%, 15.5% and 24.1% respectively of the total number of reads and 28.1%, 23.7% and 18.2% respectively of the number of reads identified as yeast species (Figure 3).

**Figure 3:**
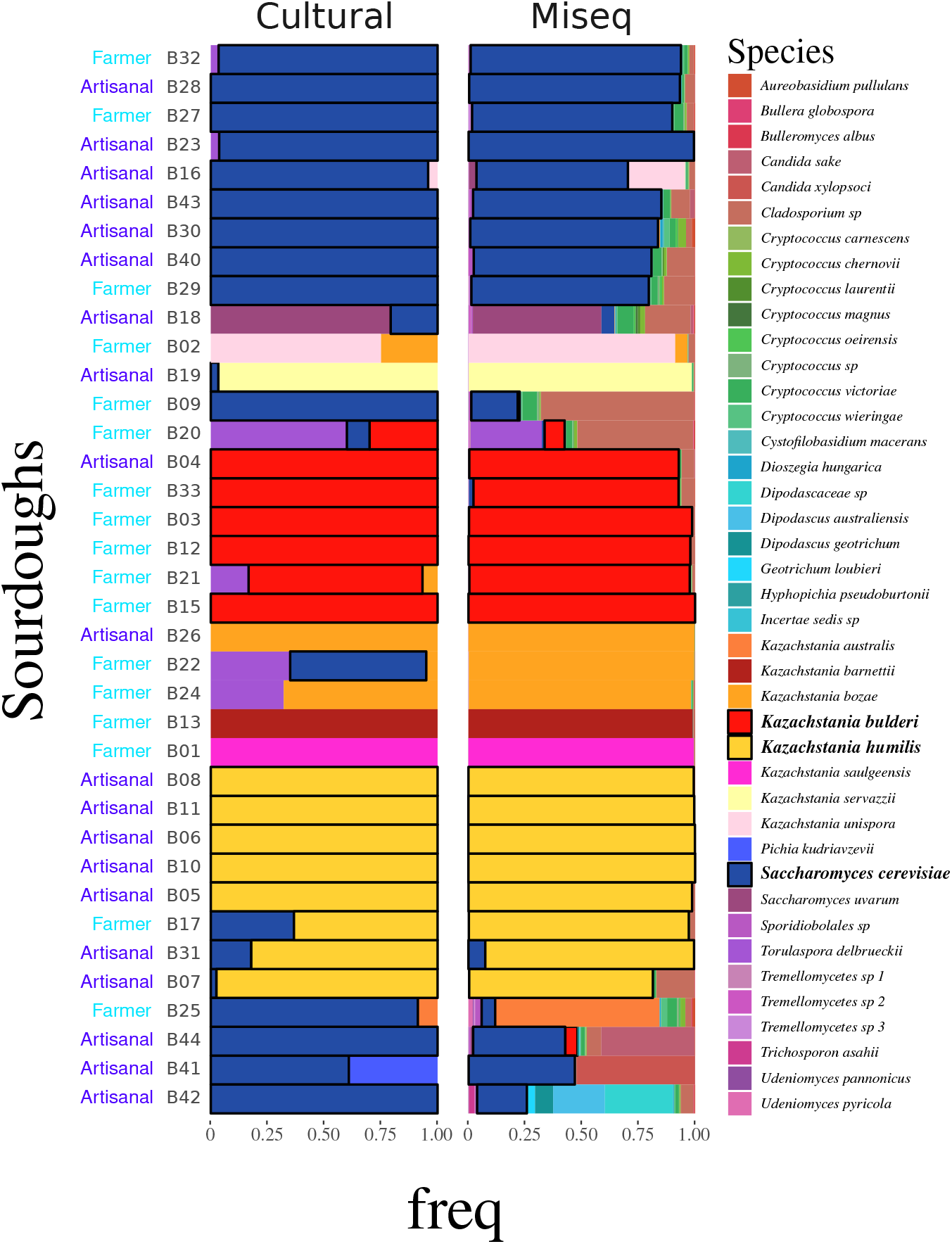
Fungal species diversity was analyzed for 38 out of the 39 sourdoughs with both cultural and metabarcoding methods. Left: species were identified by traditional microbial isolation and identification using ITS sequencing. Right: species were identified using ITS1 metabarcoding. The three most frequently-encountered species are shown in contrasting colors surrounded by black (blue: *Saccharomyces cerevisiae*, red: *Kazachstania bulderi*, yellow: *Kazachstania humilis*). The bread making practice of the baker who supplied the sourdough is indicated on the left (“artisanal” in purple, “farmer” in light blue).

The non-culture-based and culture-based methods allowed the identification of the same dominant species (defined as a species with an over 50% frequency) for all sourdoughs but five (B09, B20, B22, B25, B41) (Figure 3). In two cases, the discrepancy was explained by the detection of the *Cladosporium* genus at high frequency with metabarcoding while this species could not be isolated in the laboratory (Figure 3). In two other cases, it was explained by a high number of *S. cerevisiae* isolated in the laboratory compared to what was observed using metabarcoding sequencing. In the last case, the identification of *Pichia kudriavzevii* required additional sequencing. Because metabarcoding allows a deeper characterization of the fungal species diversity with few discrepancy cases, the distribution of fungal species across sourdoughs has been further described using metabarcoding data only (Figure 4). Previous analysis of the same sourdoughs revealed that *Fructilactobacillus sanfranciscensis* was the dominant bacterial species in all analyzed sourdoughs but two, where the dominant species was either *Latilactobacillus curvatus* or *Companilactobacillus heilongjiangensis* [20, 52, 53]. Therefore, we decided to study the species composition of the fungal community only.

**Figure 4:**
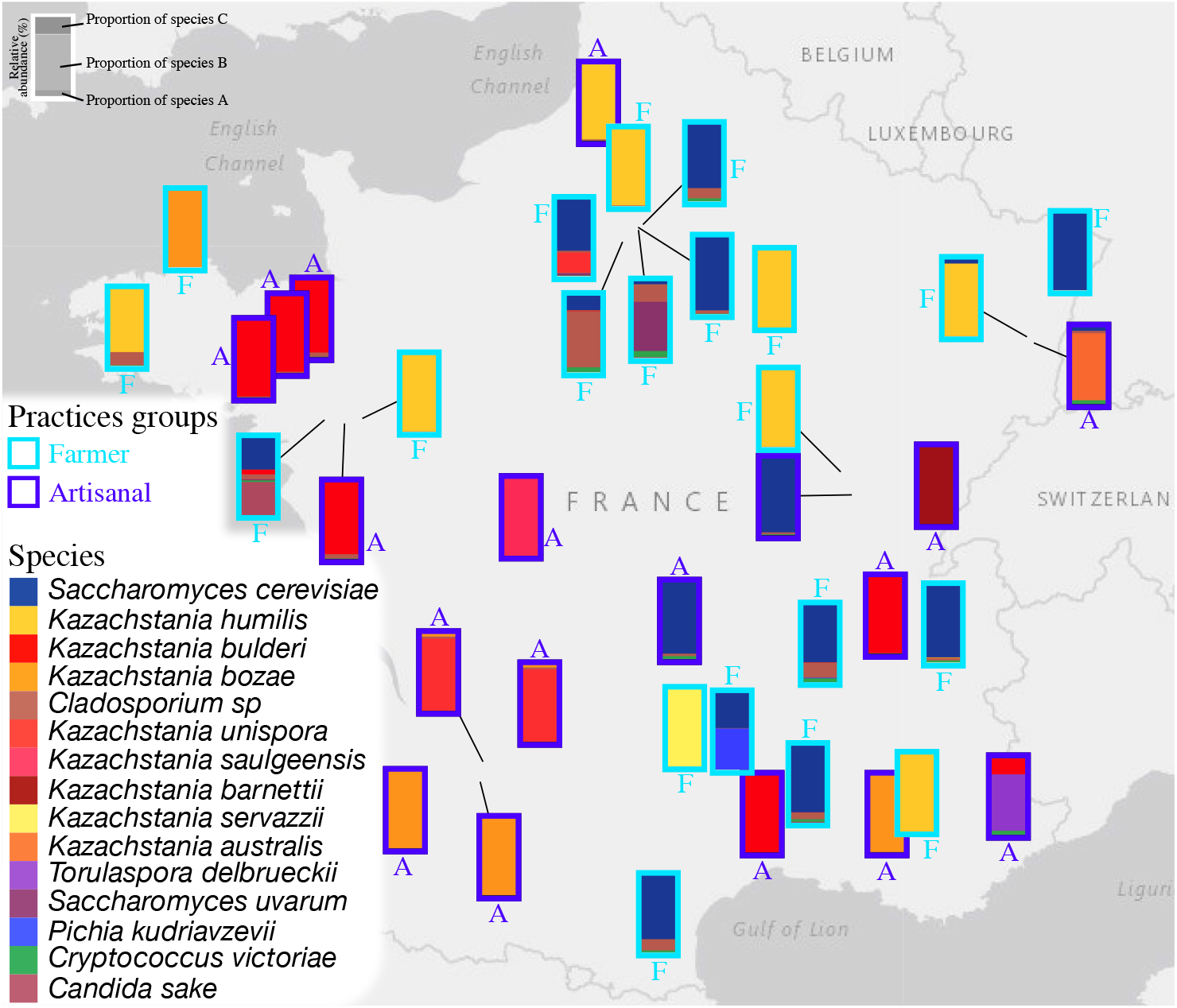
Fungal species composition of 38 French sourdoughs. Each bar represents one sourdough and is placed on the map where the sourdough was collected. Sourdoughs coming from bakers with a “farmer” practice are bordered in light blue and labelled with an “F” while sourdoughs coming from bakers with “artisanal” practice are bordered by dark blue and labelled with and “A”. Each species is represented by a different color. The relative abondance of each species, estimated by metabarcoding analysis, is represented by the area of each bar, as shown at the top left.

### Fungal species diversity within and between sourdoughs

All sourdoughs but two had a dominant yeast species with a relative abundance over 50% and many species with a lower relative abundance (Figure 3, Figure 4). Within sourdoughs, fungal species richness ranged from 10 to 33, with a 23 median (Table S3 [87]). The effective number of species per sourdough calculated from the Shannon diversity index ranged from 1 to 7 (Table S3 [87]), with 70% of sourdoughs having an index below two (Table S3 [87]). The bread making practice group (artisanal-like/farmer-like) did not influence significantly the level of fungal α-diversity in sourdough (Wilcoxon rank exact test, Wshannon = 156, p-value = 0.16, Wsimpson = 165, p-value = 0.08). Between-sourdough ß-diversity was analyzed using weighted UniFrac distances, computed from a phylogenetic tree built from the distances between OTUs using *Sporidiobolales* species as root (Figure S2 [87]). UniFrac distances computed with four differently rooted trees were highly positively correlated (Figure S3 [87]). UniFrac distances between sourdoughs ranged from 0.0005 and 0.71, with a median of 0.49 and a mean of 0.52. The clustering of sourdoughs according to their UniFrac distances is shown Figure 5A. There was no significant correlation between the UniFrac distances and geographical distances between sourdoughs (Mantel test, P=0.35).

**Figure 5:**
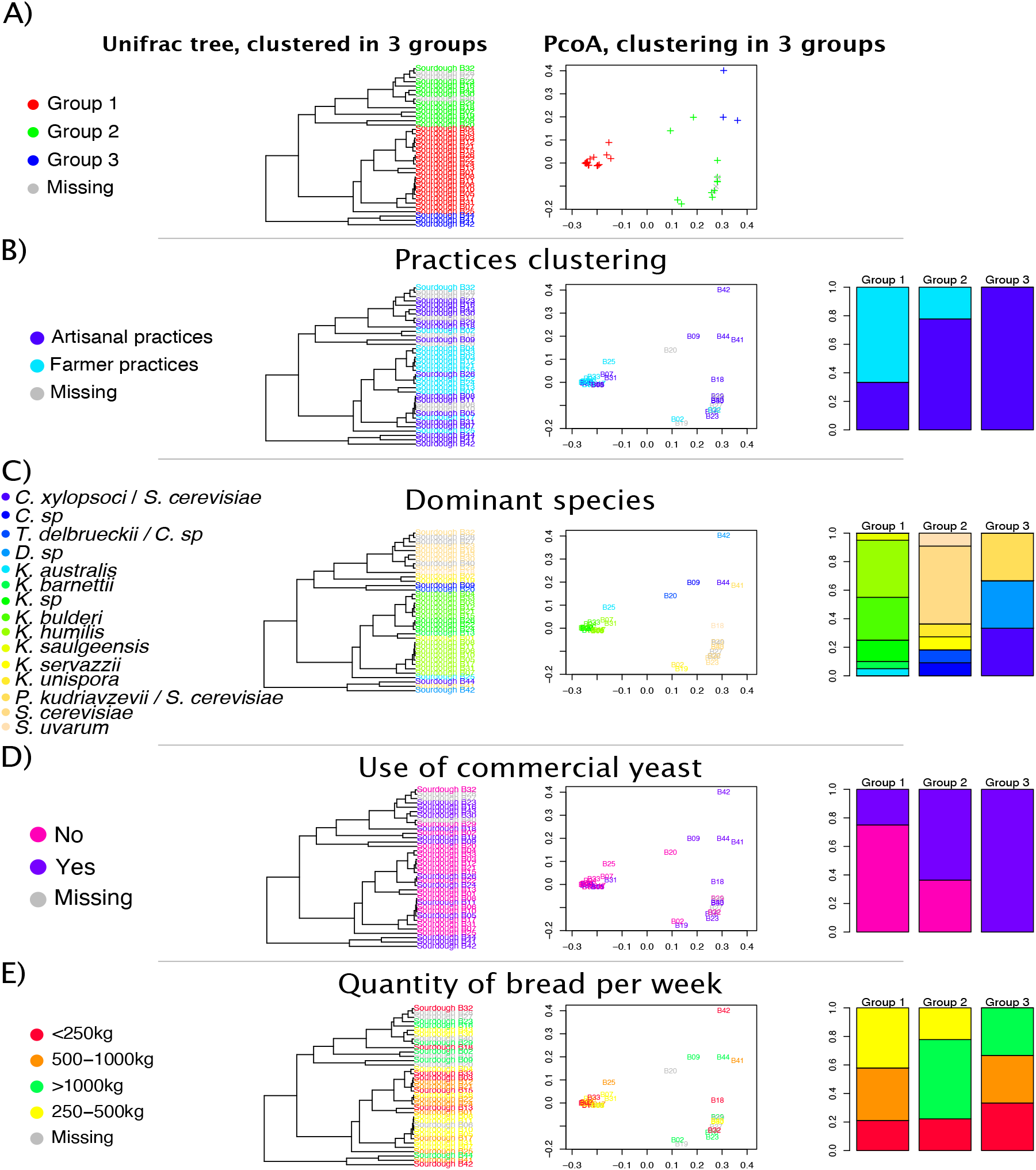
Clustering of sourdough fungal communities based on weighted UniFrac distances and association with baking practices. The weighted UniFrac distance between communities takes into account the relative abundance of the observed species as well as their phylogenetic relationships. In all panels, the clustering of sourdoughs fungal communities based on their Unifrac distances is shown on a tree on the left and on a Principal Component Analysis (PcoA) in the center. Panel A shows the three groups of fungal communities formed based on their weighted UniFrac distances. Panels B through E highlight the distribution of a principal variable in these three groups of fungal communities. The distribution is shown by coloring the different levels of the variable on the tree and on the PCoA but also as a barplot on the right side of the panel. Panel B shows the distribution of the two types of bread making practices, panel C the dominant yeast species, panel D the use of a commercial yeast starter, panel E the amount of bread made each week. Group 1 contains sourdough fungal communities whose dominant species is a *Kazachstania* species. Group 2 contains sourdough fungal communities whose dominant species is *Saccharomyces cerevisiae*. Group 3 contains sourdough fungal communities in which two yeast species co-occurred, one being *S. cerevisiae*. The yeast genus in panel C is abbreviated with, *C*. for *Candida, D*. for *Dipodascaceae, K*. for *Kazachstania, S*. for *Saccharomyces, T*. for *Torulaspora*.

We then analyzed specifically the occurrence of yeast species in sourdoughs as yeasts, together with lactic acid bacteria, are the main functional players in a sourdough ecosystem and for bread quality (Figure 4). Over the 40 yeast species detected in the 38 sourdoughs, 12 had a relative abundance over 50% in at least one sourdough, four had a relative abundance between 20% and 50% and 24 had a relative abundance below 10%. All dominant species (relative abundance over 50%) were fermentative yeast species, except in one sourdough that had a *Cladosporium* species as co-dominant species. We found all the sourdough yeast genera (*Saccharomyces, Kazachstania, Pichia, Torulaspora* and *Hyphopichia*) commonly reported in the literature except the *Wickerhamomyces* genus that we did not detect in our samples [1, 15, 16].

### The baker’s yeast species, *Saccharomyces cerevisiae* is not the most widespread yeast species in French organic sourdoughs

*Saccharomyces cerevisiae* was found in 53% of all sourdoughs (95% confidence intervals=36% - 69%) but was dominant (relative abundance over 50%) in only 24% (95% confidence intervals=11% - 40%) (Figure 3, Figure 4). In two cases, *S. cerevisiae* co-occurred with another yeast species at similar relative abundance. In the first case, *S. cerevisiae* was present at a relative abundance of 40% with *Candida sake* at a 41% relative abundance. In the second case, it was found at a relative abundance of 47% with *Pichia kudriavzevii* at a relative abundance of 52%. In all the other cases, *S. cerevisiae* had a relative abundance below 21% and was found with other dominant yeast species, such as *Kazachstania australis, Kazachstania humilis, Saccharomyces uvarum* or *Torulaspora delbrueckii*. This suggests that *S. cerevisiae* did not displace other species and can indeed be out-competed by other species in sourdoughs.

### Sourdough yeast species mostly belong to the *Kazachstania* genus

*Kazachstania* was the most represented yeast genus over all sourdoughs, when considering both the number of reads over all sourdoughs and the number of detected species. This genus represented 57% of the total number of reads while *Saccharomyces* represented 26% of the total number of reads. In addition, eight species of the *Kazachstania* genus were found in sourdough, while the *Saccharomyces* genus was represented by two species (*S. uvarum* and *S. cerevisiae*) (Figure 3, Figure 4). The *Kazachstania* genus is one of the closest genetically related genus to *Saccharomyces* and contained Crabtree positive yeasts, able to ferment glucose even when oxygen is present if the amount of sugar is sufficient (Hagman & Piskur 2015). *Kazachstania* species dominated in 54% (95% confidence intervals=36%-69%) of sourdoughs while *Saccharomyces* species dominated in 27% only (95% confidence intervals=13%-43%). *Kazachstania humilis*,followed by *K. bulderi*, were the most commonly dominant *Kazachstania* species, and found in respectively 21% (95% confidence intervals=10%-37%) and 15% of sourdoughs (95% confidence intervals=6%-31%). A recently described *Kazachstania* species, *Kazachstania bozae*, was also identified in five sourdoughs (4.5%- 29%) and found dominant in three (1.7%-22%) [64]. Strains of this species were closely related to a strain previously isolated from boza, a Bulgarian fermented drink, as estimated with ITS and LSU (D1D2) barcodes (Source: NCBI, GenBank: KC118125.1 and KX369579.1). In addition, *Kazachstania saulgeensis*, a recently described species [65, 66], was dominant in one sourdough (0.07%-14%). *Kazachstania unispora* and *Kazachstania servazzii* which had previously been detected in sourdough were also found [17, 17, 53, 57, 58, 63, 67–71]. Some *Kazachstania* species were detected for the first time as dominant in sourdoughs, whereas they had been previously found in other environments, like soil (*K. australis*) and sauerkraut (*K. barnettii*) [72–74]. None of the previous studies on sourdough have observed as many *Kazachstania* species in sourdough.

### The composition of sourdough fungal communities was associated with differences in bread making practices

We tested whether sourdough fungal community beta diversity could be explained by bread making practices. To do so, we performed univariate PERMANOVA analysis on the 30 bakers with fewer than 8 missing values for the 29 bread making practices variables (Table S4 [87]). The univariate analysis revealed that the weighted UniFrac distance was structured according to the use of commercial yeast in bakery (P<0.05). It also varied significantly with sourdough age, chief sourdough origin (dough, sourdough or both), the quantity of bread produced per week, the milling method (cylinder, millstone, Astrie, Tyrol), the type of wheat variety (ancient, modern or a mix thereof) and the fermentation duration. However, after FDR correction for taking into account multiple testing, none of these variables significantly explained UniFrac distances.

In order to understand further the relationship between sourdough fungal community composition and bread making practices, we clustered sourdoughs according to their fungal community composition, on the basis of the PCoA of their weighted UniFrac distances. Then, we tested the link between the fungal community group and the bread making practice group (farmer/artisanal practice group) as well as the link between the fungal community group and each of the different bread making practices (Figure 5).

Sourdoughs were clustered into three fungal community groups. Group 1 encompassed all sourdoughs (but two) having *Kazachstania* species as dominant species (*K. humilis, K. barnettii, K. bulderi, K. saulgeensis, K. bozae*). Group 2 contained sourdoughs with *Saccharomyces sp., K. servazzii* or *K. unispora* as dominant species. Group 3 harbored sourdoughs with *S. cerevisiae* together with other species such as *Pichia kudriavzevii, Candida sake*, or a Dipodascaceae sp. Group 1 sourdoughs were mostly made by bakers having farmer’s bread making practices while group 2 and group 3 sourdoughs were mostly made by bakers using artisanal practices (exact Fisher test, P=0.035). The fungal community groups were significantly associated with two specific bread making practice variables: the quantity (in kg) of bread made per week (Exact Fisher test, P=0.001) and the use of commercial yeast (Exact Fisher test, P=0.05). All sourdoughs in group 2 but one were found in bakeries making between 500 kg and 1000 kg of bread per week, while groups 1 and 3 sourdoughs originated from bakeries producing very different amounts of bread (ranging from amounts below 250 kg to over 1000 kg). In addition, group 1 sourdoughs were more frequently found in bakeries that do not use commercial yeast while groups 2 and 3 were more frequently found in bakeries using the commercial yeast *S. cerevisiae* (Exact Fisher test, P=0.01). Interestingly, group 1 sourdoughs harbored *S. cerevisiae* either at a relative abundancy below 1% or not at all, while all groups 2 and 3 sourdoughs had *S. cerevisiae* at a relative abundancy over 20%, except in three cases where it was either absent or at a relative abundancy below 6%.

To test more specifically the link between bread making practices and the distribution of *Kazachstania* species, we analyzed more in-depth group 1 sourdoughs. Within this group, eight sourdoughs had *K. humilis* as dominant species, six had *K. bulderi*, three had *K. bozae* and the remainder had still other *Kazachstania* species. All sourdoughs made with artisanal practices carried *K. humilis* as dominant species or, in one case, the *K. bozae*. By contrast, sourdoughs made with farmer practices had as dominant species *K. bulderi, K. australis, K. barnettii, K. saulgeensis* or *K. bozae* (exact Fisher test, P=0.004).

### Fungal community composition was partly related to sourdough acidity, maltose concentration and hydration

The composition of fungal community may affect sourdough metabolic content (sugars, acids, alcohols) via fungal strains metabolite consumption and production. Inversely, the presence and concentration of different compounds (sugars, acids, alcohols) may affect differently the fitness depending on the strains and consequently be one of the drivers of fungal community composition. For example, lactic acid bacteria (LAB) are the main producers of acidity in sourdough, but yeasts also produce acetic acid and also indirectly affect acidity through positive or negative interaction with bacteria.

To investigate the relation between sourdough fungal communities and metabolic compounds, we quantified sourdough hydration, yeast density, bacteria density, sourdough pH, total titrable acidity (TTA), sourdough concentration in seven sugars (maltose, glucose, fructose, raffinose, arabinose, mannose, xylose), four alcohols (glycerol, ethanol, mannitol, meso-erythtritol), six acids (lactate, acetate, glutarate, pyruvate, malate, succinate) and calculated the fermentative quotient (lactate over acetate ratio). For each variable, there was a wide range of variation (Table S5 [87]). The principal component analysis based on all variables showed no evidence of sourdough grouping (Figure S4 [87]). As expected in fermentation, yeast density was positively correlated to ethanol (r=0.74, P<0.001), glycerol (r=0.67, P<0.001), and acetate (r=0.6, P<0.001) concentration. However, it was not significantly correlated to sugar concentrations. This might be explained by the co-occurrence of bacteria which have their own metabolism and interact by competition and/or cross feeding with sourdough yeasts.

We then tested whether the variation of each quantitative variable was associated with the bread making practice groups (farmer-like practices and artisanal practices). There was no significant effect of the bread making practice group except for sourdough hydration that was significantly higher in sourdoughs made using farmer-like practices (F1,94=11,69, P<0.001). On average, sourdoughs made with farmer-like practices had 55% water while sourdoughs made with artisanal-like practices had in average 49% of water.

In addition, we tested whether variations in quantitative variables were associated with the fungal community groups (Table S5 [87]). Group 3 microbial community sourdoughs (defined by PCoA clustering on UniFrac distance, see below), which contains *S. cerevisiae* in co-dominance with a second yeast species (*Candida sake, Pichia kudriavzevii* or a *Dipodascus* species), had a significantly higher mean pH (mean pH_group3_=4.2 against pH _group1_=3.8, Tukey Contrasts, P<0.001), lower TTA (mean TTA _group3_=7.7 against TTA _group1_=17.1, Tukey Contrasts, P=0.002), and a higher maltose concentration (mean Maltose _group3_=52.8 mg/gr of sourdough against Maltose _group1_=24.1 mg/gr of sourdough, Tukey Contrasts P=0.002) than group1, having a *Kazachstania* dominant species. Compared to group 2 having in most cases *S. cerevisiae* as dominant species, it also had higher pH (pH _group2_=3.9, Tukey Contrasts, P=0.003), and higher maltose concentration (Maltose _group2_=23.7, Tukey contrast, P=0.003). These data may reflect a lower fermentative activity for group 3 fungal community having two co-dominant species, and/or a negative interaction effect of group 3 fungal community on the activity of lactic acid bacteria (LAB), which are the main producers of sourdough acids. Previous studies on the bacteria content of the same sourdoughs showed that *F. sanfranciscensis* was most generally the dominant species, although *C. heilongjiangensis, L. curvatus* or *Levilactobacillus brevis* were also found as dominant species [20, 52, 53]. We found no significant correlation between LAB and yeast densities (r= −0.15, p= 0.45, Figure S4 [87]) but the link between fungal and bacterial community might be species and strains dependent. Additional studies on the interactions between fungal and bacterial communities need to be performed to better understand how they may drive sourdough acidity and sugar content.

We also analyzed whether the variations of each quantitative variable was associated with the dominant yeast species. We only considered the 26 sourdoughs having either *S. cerevisiae* (9 sourdoughs), *K. humilis* (8 sourdoughs), *K. bulderi* (6 sourdoughs) or *K. bozae* (3 sourdoughs) as dominant species, since the other yeast species were found dominant only once. The differences in dominant species was not significantly associated to variation in sourdough sugar, acid or alcohol concentrations. However, on average, sourdoughs dominated by *K. bulderi* were more hydrated (63% water content in average) than sourdoughs dominated by *K. humilis, K. bozae*, and *S. cerevisiae*, having respectively 49%, 47%, 53 % water content in average (P<0.001 for the 3 Tukey Contrasts). *Kazachstania bulderi* was found to be dominant only in sourdoughs made using farmer-like practices, a bread making practice group that was also found to be associated with more hydrated sourdoughs. Additional experiments should be carried out to test whether this species has indeed a better fitness in more hydrated sourdoughs or whether its presence in more hydrated sourdoughs is related to covariation with other farmer practices.

In conclusion, no clear evidence was found of the impact of bread making practices or of the dominant yeast species on the metabolic composition of sourdough. On the other hand, our results showed metabolic differences between sourdoughs having one or two co-dominant yeast species.

### Phenotypic signatures of domestication

A previous analysis on *S. cerevisiae* revealed that sourdough strains had higher average fitness and fermentation performance than strains from other environments in a sourdough-mimicking medium [10]. Here, we investigated whether evidence of a domestication syndrome could also be found in *K. humilis* and *K. bulderi*, the two *Kazachstania* species most commonly found in French sourdoughs. We tested whether fitness (log of population size and mortality at the end of fermentation) and fermentation performance (CO_2_max, Vmax, t1g, tVmax) differed between sourdough strains and strains from elsewhere.

A principal component analysis of 38 strains of *K. bulderi* and *K. humilis*, based on quantitative variation in the six phenotypic variables described below was carried out. The first two axis explained 80.5% of the variation and clearly separated strains by species (Figure 6). The *K. bulderi* strains were located at the right of the PCA and were characterized by high population size and low mortality at the end of fermentation, while the *K. humilis* strains were located at the left and were characterized by a rapid onset of fermentation (t1g), high maximum fermentation rate (Vmax), and a short time to reach Vmax (tVmax). Non-sourdough strains of *K. humilis* were located outside the cloud of sourdough strains while non-sourdough strains of *K. bulderi* were distributed within and outside the cloud of sourdough strains.

**Figure 6:**
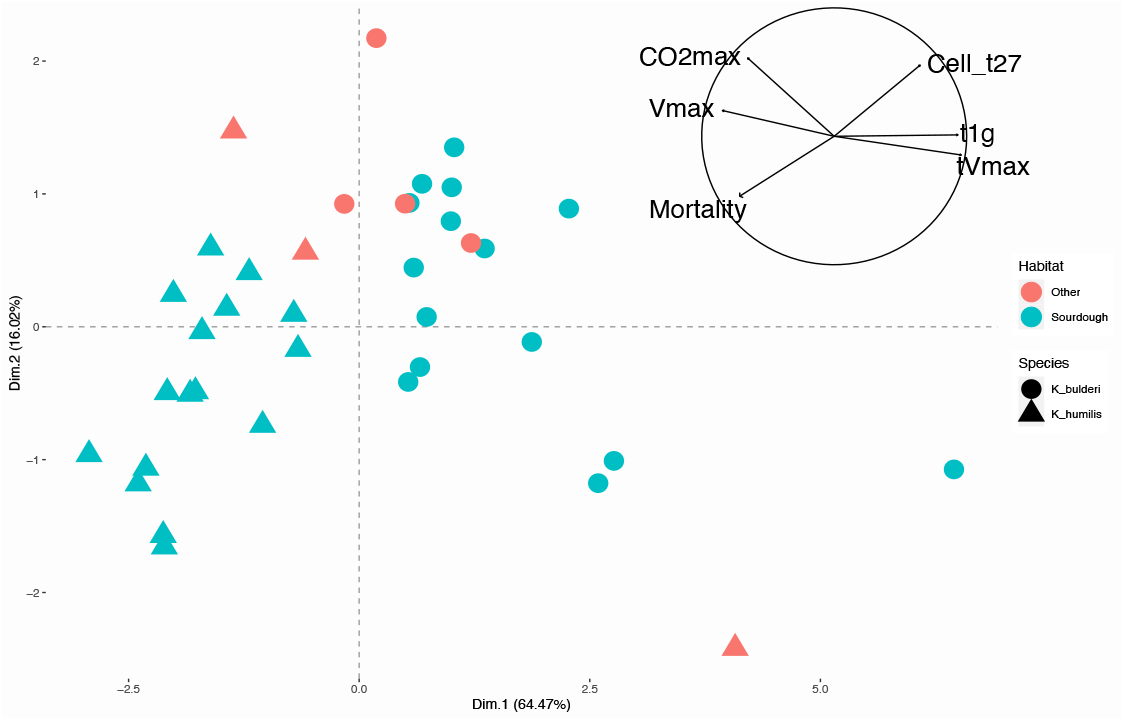
Principal component analysis of 37 *K. humilis* and *K. bulderi* strains based on the quantitative variation of maximum CO_2_ production (CO_2_max), fermentation latency phase (t1g), maximum CO_2_ production rate (Vmax), time to reach the maximum production rate (tVmax), log of population size and mortality at the end of the fermentation. The correlations between variables are presented on the left while the figure on the right shows the projection of strains on the first two axes representing 70.64% of the variation. The strains are colored according to their habitat of origin. Sourdough strains are indicated in blue and non-sourdough strains in red. Their species is indicated by symbol. *Kazachstania humilis* strains are indicated by a circle and *Kazachstania bulderi* strains by a triangle.

Statistical comparisons of sourdough and non-sourdough strains of *K. bulderi* and *K. humilis* for each phenotypic variable revealed phenotypic divergence for *K. humilis* but not for *K. bulderi*. While the *K. bulderi* sourdough strains did not ferment significantly faster than the non-sourdough strains, the *K. humilis* sourdough strains showed significantly higher Vmax, lower t1g, and lower tVmax than the non-sourdough *K. humilis* strains (Figure 7, Table S6 [87]). On average, they started fermentation two hours before the others and reached Vmax three hours before the others. In addition, their Vmax were on average 34% higher than the others.

**Figure 7:**
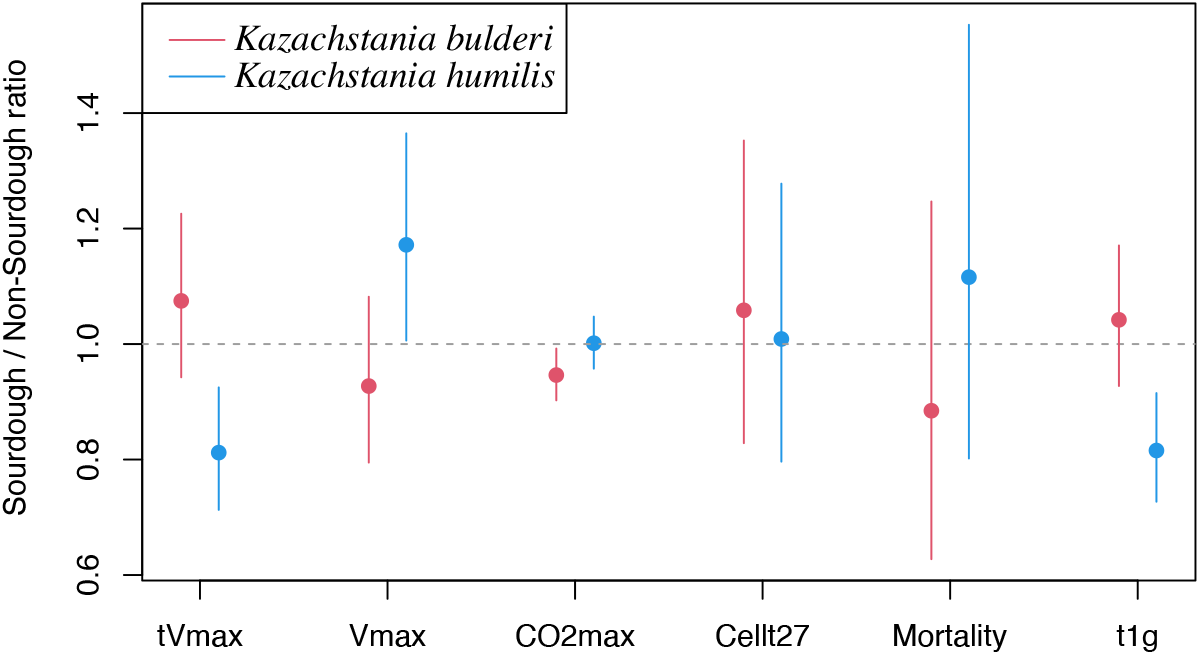
Ratio between the sourdough strains mean and the non-sourdough strains mean values of each quantitative variable measuring fermentation performance: time to reach the maximum production rate (tVmax), maximum CO_2_ production rate (Vmax), maximum CO_2_ production (CO_2_max), log of population size after 27 hours of fermentation (Cellt27), cell mortality after 27 hours of fermentation (Mortality), fermentation latency phase (t1g). Confidence intervals are indicated by bars. Mean ratio and confidence intervals are shown in red for *Kazachstania bulderi* and blue for *Kazachtania humilis*.

## Discussion

Sourdough microbial diversity has been intensively studied worldwide. Despite a cultural and historical interest on bread in France, French sourdough fungal diversity was only partly characterized before this study [19, 53, 71]. A recent large-scale (> 500 starters) study of sourdough microbial diversity revealed the fungal diversity that can be detected over the globe across home-made sourdoughs [17]. All the yeast species detected at a relative abundance over 1% in this international collection of sourdoughs were detected in French sourdoughs except the species *Wickerhamomyces anomalus, Pichia membranifaciens, Naumovozyma castellii* and *Saccharomyces bayanus*. Inversely, French baker’s sourdough harbored some yeast species that were never found elsewhere, such as *K. bozae, K. australis*, and *K. saulgeensis* [65].

Beyond the genus of the baker yeast species, *S. cerevisiae*, the most represented yeast genus in French sourdough was *Kazachstania*. Eight *Kazachstania* species were found as dominant yeast species in at least one French sourdough: *K. humilis, K. bulderi, K. barnettii, K. unispora, K. servazzii, K. bozae, K. australis* and *K. saulgeensis*. Three *Kazachstania* species (*K. exigua, K. lodderae, K. naganishii*) already reported in sourdough were not found in our collection of French sourdough. *Kazachstania lodderae* and *K. naganishi* are rarely found in sourdough. By contrast, *K. exigua* is a frequently cited sourdough species in the literature. This species has been previously found in France, Finland [77], Italy [78, 79], Denmark [80], Ethiopia [81], USA [17] and is the first species to have been isolated from a sourdough (in San Francisco, [29]). However, its taxonomic characterization may have been hampered by the fact that it probably originated from hybridization between unknown yeast species [82]. To date, the genus *Kazachstania* is composed of more than 40 species, of which 11 are present in sourdough. It is possible that an adaptive radiation linked to the adaptation to different sourdoughs or to different anthropized niches has taken place, as it has been observed for example in cichlids during their adaptation to different lakes or in *Penicillium* domesticated fungi. Indeed, five of the *Kazachstania* species present in the sourdough have so far only been detected in human-related niches. These are *K. saulgeensis, K. barnettii, K. bozae, K. bulderi* and *K. humilis*. These species are genetically closer to each other than they are to *K. servazzii* and *K. unispora* which have also been found in nature and are grouped in another part of the *Kazachstania* pylogenetic tree. Genomic analysis of these species would shed light on their evolution and the genetic changes that would have been selected during potential domestication. So far, eight of the 11 *Kazachstania* species found in the sourdough have at least one genome assembly available in public databases, including *K. saulgeensis* and *K. barnettii* [44, 45], and the assemblies of *K. bulderi* and *K. humilis*, which were recently published in public databases (Bio project: PRJEB44438). The genomic and phenomic analysis of the large collection of *Kazachstania* strains obtained by our study, together with the world collection of *Kazachstania* strains, may shed light on the radiation and domestication processes of these species.

We found that yeast community composition was partly related with bread making practices. Bread making practice divergence also led to different phenotypic signatures. Strains of the *K. humilis* species, which was typical of sourdough made by artisanal-like practices, had higher fermentation rate while strains of *K. bulderi* which was typical of sourdough made by farmer-like bread making practices had not. The species *K. humilis* has been found in many countries, viz. Austria, Canada, China, Denmark, Ethiopia, Finland, Germany, Greece, Italy, Morocco, the Netherland, Spain, UK, USA, and France [1, 15–17, 19, 53, 55–60]. It is also the most frequently encountered *Kazachstania* species in sourdoughs around the world. This species is therefore frequently found in bakeries, where short fermentations are often favored. This may explain why sourdough strains of *K. humilis* seem to have been selected for increased fermentation rate. Increased fermentation rate was also found in bakery strains of *S. cerevisiae* when compared to non-bakery strains.

In contrast, we did not find evidence for improved fermentation performance in *K. bulderi*, which was the third most represented species in French sourdough. *Kazachstania bulderi* was found in bakeries with farmer-like practices. These bakeries often bake bread once or twice a week and store their sourdough for several days. They also often use long fermentation and thus may not have selected an increased fermentation rate. Farmer-bakers typically store their sourdough for several days and therefore make a lower number of back-slopping. In addition, they make bread with longer fermentation times than artisanal bakers. It is therefore possible that they did not select to accelerate the speed of fermentation and instead let natural selection in the sourdough environment act alone. Alternatively, the lack of phenotypic divergence between sourdough and non-sourdough strains may reflect the limitation of our sampling. *Kazachstania bulderi* has been reported for the first time in anaerobic maize silage in the Netherlands and in fermented liquid feed for piglets [61, 62], more recently in French, Belgium and Spain sourdoughs [19, 53, 57, 63] but to our knowledge was never found in wild environment. Here, we compare sourdough strains with strains coming from silage and animal feed. It is unknown whether silage and animal feed strains are wild strains or feral strains that have escaped from other domesticated environments. This may explain why we did not detect any phenotypic divergence between sourdough and non-sourdough strains of *K. bulderi*. Other than fermentation phenotypes, there was no evidence of fitness differences between sourdough and non-sourdough strains of *K. humilis* and *K. bulderi* in sourdough mimicking media. Additional experiments in real dough should be performed to further test the effect of natural selection in this environment.

Other evolutionary processes than selection could also explain the distribution of yeast species across sourdoughs. Interviews with the bakers working with sourdough hosting *K. bulderi* and *K. bozae* suggested the role of dispersion of these species in French sourdoughs. Indeed, these bakers have been connected over the years either through seed exchanges, sourdough mixing or gifts, bread making training in common or working in one another’s bakery. Some yeast species have been found in the bakery house environment and baker’s hands and may thus be dispersed through baker’s tools or baker’s travels [58, 67, 75, 76]. However, it is still unclear whether wheat seeds and flour are a source of sourdough yeasts.

To our knowledge, this is the first evidence of the influence of artisanal practices on taxonomic diversity in microbial communities. On the other hand, several studies have shown that making fermented products could lead to the selection of divergent phenotypes and genotypes. This is the case of sourdough and industrial populations of the baker’s yeast *S. cerevisiae* that diverge from each other and have a better fermentation performance than non-baker’s strains. As for beer populations, sourdough populations have acquired a better capacity to assimilate maltose, linked at least in part to an increase in the number of copies of the genes involved in the assimilation of maltose. Several studies on wine populations of *S. cerevisiae, Torulaspora delbrueckii* and *Lachancea thermotolerans* also showed that wine populations are genetically differentiated from strains from other environments and present beneficial phenotypes in grape must and for wine quality. The analysis of the filamentous cheese fungi *P. roqueforti* and *P. camemberti* also revealed genetically differentiated cheese populations. Interestingly, different genetic groups associated with different cheese making practices were found. Strains of the blue cheese fungus, *P. roqueforti*, isolated from Roquefort cheeses were more diverse and were genetically and phenotypically different than strains used to make other blue cheeses [83, 84, 85]. Two varieties of the soft-cheese making fungus, *P. camemberti*, with different phenotypic features, were associated with different kinds of cheese (Camembert and Brie). All together these studies show that the diversity of practices used to make fermented products allows to maintain genetic, phenotypic and taxonomic diversity.

However, fungal domestication also involved strong bottlenecks. For example, the low level of genetic diversity found in blue-cheese *P. roqueforti* strains and soft-cheese *P. camemberti* strains revealed the risk of diversity erosion in organisms used for fermented product making [6, 7, 84]. This risk is accentuated by fertility depression among fungal domesticated strains [86]. This risk is also associated with the massive use of few industrial strains or the need to standardize products or meet the specifications of industrial production or protected designation of origin (PDO) [6]. Here, we show that despite the recurrent use of *S. cerevisiae* as industrial starter species in bakeries and homes, and the occurrence of this species in a wide range of habitats such as soil, trees, and humans [9, 54], this species does not appear to have overwhelmingly colonized French traditional sourdoughs (Figure 4). This result confirmed a recent analysis which revealed that *S. cerevisiae* sourdough strains had a different evolutionary history from industrial strains [10]. The dynamic of microbial species colonization and invasion in food environment remains largely unknown. Additional experiments at the level of microbial community will shed light on the dynamic of microbial community establishment in food production and on the ability of industrial strains to invade food microbial community.

In conclusion, a great diversity of bread making practices and fungal community composition was found in our sample of French sourdoughs. Surprisingly, the well-known baker’s yeast Saccharomyces cerevisiae was found dominant only in one fourth of the sampled sourdoughs. By contrast, several species of the neighboring genus *Kazachstania* were detected at high frequency, revealing a major role for this mostly unknown genus in the study of fungal domestication and in bread making. Therefore, our results highlight the necessity of maintaining socio-cultural diversity to maintain microbial diversity in food systems. These findings could not have been revealed without the collaboration of bakers and scientists, showing the importance of participatory research projects to gain new insights into biodiversity preservation.

## Acknowledgements

We thank all the bakers and farmer-bakers for constructing the project with us and for sharing their sourdoughs and knowledges. We thank Yoann Robert, Candice Aulard, Josette Bessière for lab assistance and Matthieu Barret for MiSeq sequencing. We also would like to thank Philippe Roussel for sharing his knowledge about French bread. Preprint version 6 of this article has been peer-reviewed and recommended by Peer Community In Evolutionary Biology (https://doi.org/10.24072/pci.evolbiol.100154).

## Data, scripts, code, and supplementary information availability

Data, scripts, code and supplementary information are available online: DOI 10.5281/zenodo.5849058

