## Supplemental M&M, figures, tables for "Artisanal and farmer bread making practices differently shape fungal species community composition in French sourdoughs"

**This PDF file includes:**

Materiels and methods supplementary information
Fig. S1 to S4
Table S1 to S5

**Other supplementary materials for this manuscript include the following:**

Michel Elisa, Masson Estelle, Bubendorff Sandrine … Sicard Delphine (2022). Artisanal and farmers bread-making practices differently shape fungal species community diversity in French sourdoughs (Version 3) [Data set]. Zenodo. http://doi.org/10.5281/zenodo.5849058

### Material and methods supplementary information

#### Method S1

For each sourdough, three independent replicates of 1 gr were clarified by Carrez reagents I and II (250 µL) and centrifuged at 13.000 rpm for 5 min. The supernatant (water-soluble extract) were then frozen (- 20°C) waiting for the preparation procedure before HPLC analysis. HPLC analysis were performed with a HP 1100 LC system (Agilent technologies, Santa clara, CA, USA) equipped with a refractive index detector (RID Agilent G1382A) and a UV detector (Agilent G1314A).

Two alcohol (ethanol, glycerol), six organic acids (Lactacte , acetate, glutarate, pyruvate, malate, succinate), and two sugar (Arabinose, Meso-Erythritol) were quantified on a Rezex ROA-organic acids column (SDVB - $H^+$ 8%, 300x7.8mm, Phenomenex, Torrance, CA, USA) using the following conditions : mobile phase 0.005N H2SO4, flow rate 0.6 mL/mn, column temperature 60°C, injection volume 15µL, run time 25 mn. The extracts were diluted 6-fold with the mobile phase before injection in order to obtain an acidic sample. The organic acids were quantified thanks to a UV detection (wavelenght of 210 nm) while the sugars and alcohols were measured with a RID detector (temperature 35°C).

One alcohol (sorbitol) and six sugars (maltose, glucose, fructose, raffinose, mannitol, xylose, ribose), which could not be well separated by the first column were quantified using a Rezex RPM-monosaccharide column (SDVB – Pb+2 8%, 300x7.8mm, Phenomenex, Torrance, CA, USA) using the following conditions: eluant ultrapure water, flow rate 0.6mL/mn, column temperature 80°C, RID detector temperature 40°C, injection volume 12µL, run time 45 mn. The sourdough extracts were diluted with the mobile phase 0.005N H2SO4, but in a larger proportion than for the Rezex ROA-organic acids column to be close to neutral pH (9 volumes of water for 1 volume of extract). All concentrations were calculated by external calibration using HPLC or analytical grade standards, purchased from SIGMA. Each of them was injected at 3 or 4 concentrations levels chosen in accordance with the expected concentration, from preliminary sourdough analysis. Metabolite concentrations were corrected by the density of each sample due to sugar content.

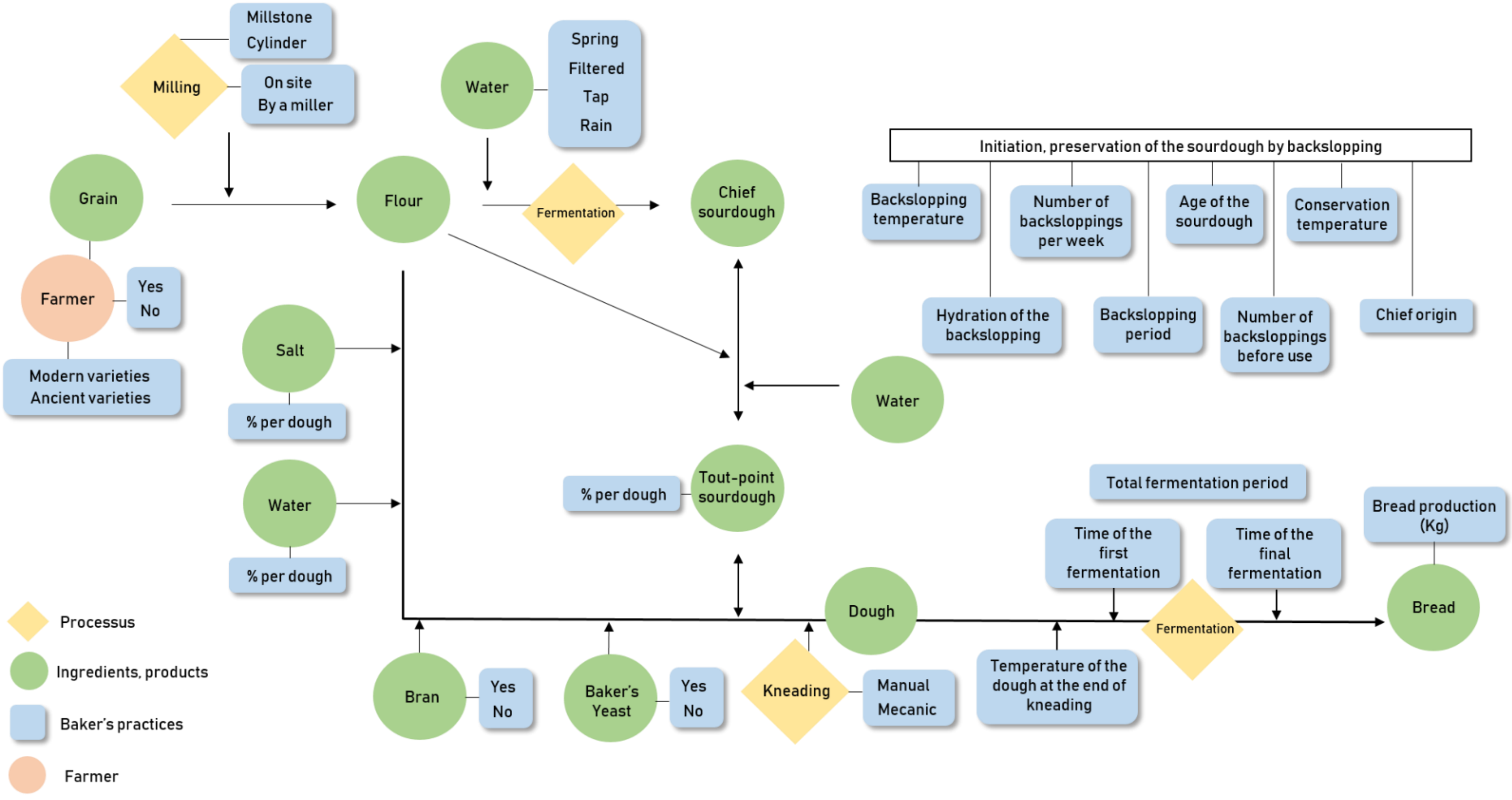


**Fig. S1.** Descriptive scheme of the survey on bread-making process. In yellow are indicated technological process, in green ingredients used in bakery chain to make bread, in blue the practices variables that were reported from bakers and in orange farmers.

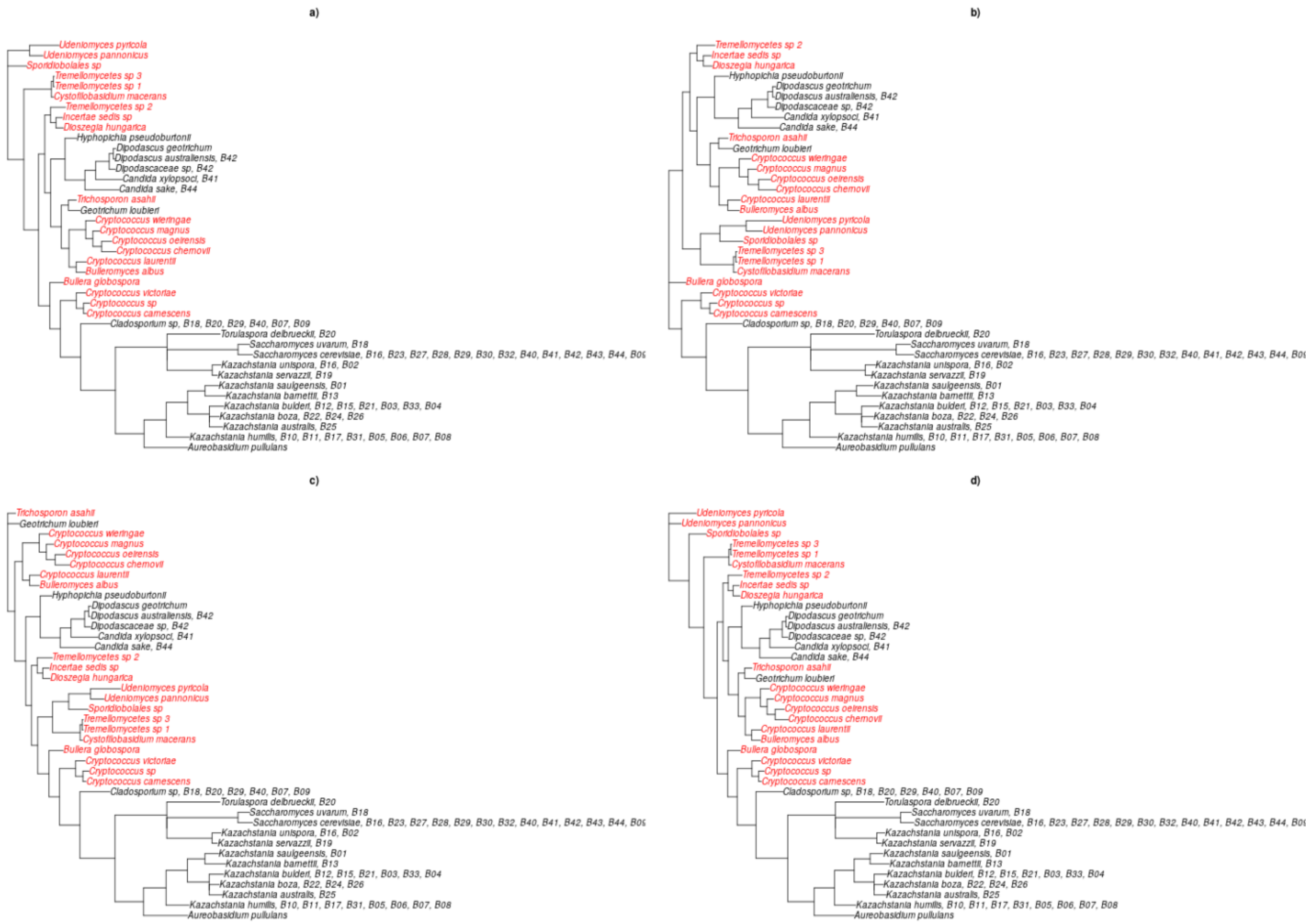


**Fig. S2. Trees made from OTUs genetic distances with different roots.** Unifrac distances were computed from rooted phylogenetic trees built from the distances between OTUs. When several OTUs were identified as the same species, we kept the most representative sequence. We investigated the impact of the chosen root on the phylogenetic tree and on the computed unifrac distances. The results presented in the main text were obtained using the phylogenetic tree rooted on the OTU identified as *Sporidiobolales species*. Here, we compared these results with those obtained from trees rooted on *Bullera globospora*, *Udeniomyces pyricola* and *Trichosporon asahii*. Rooted trees on four different OTU identified as: a) *Sporidiobolales species*, b) *Bullera globospora*, c) *Trichosporon asahii*, d) *Udeniomyces pyricola*. Species name are colored according to their phylum. Baker's codes are shown in front of each species when their frequency were over 10 % in the sourdough.

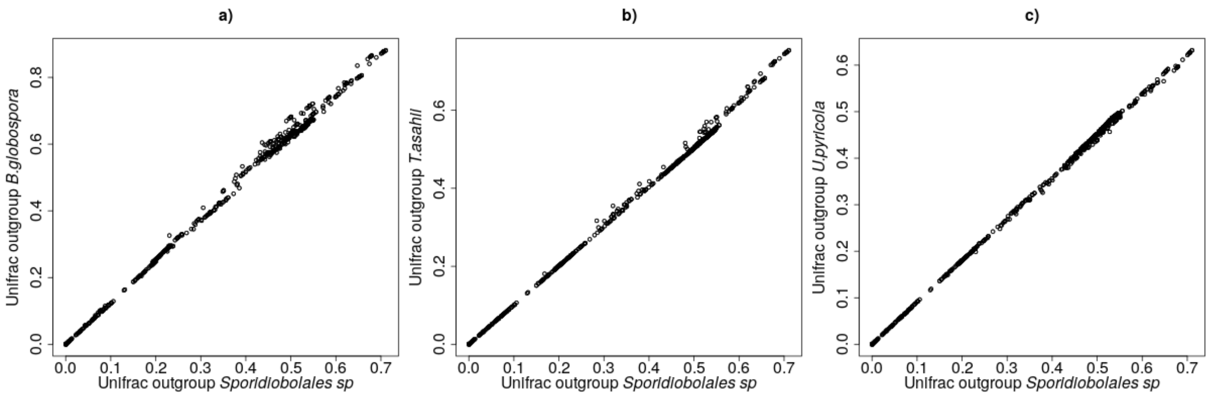


**Fig. S3.** Unifrac distances computed with rooted trees on four different species. On the x-axis the Unifrac distance computed with the tree rooted on Sporidiobolales species. On the y-axis, Unifrac distance computed with the tree rooted on: a) *Bullera globospora*, b) *Trichosporon asahii*, c) *Udeniomyces pyricola*.

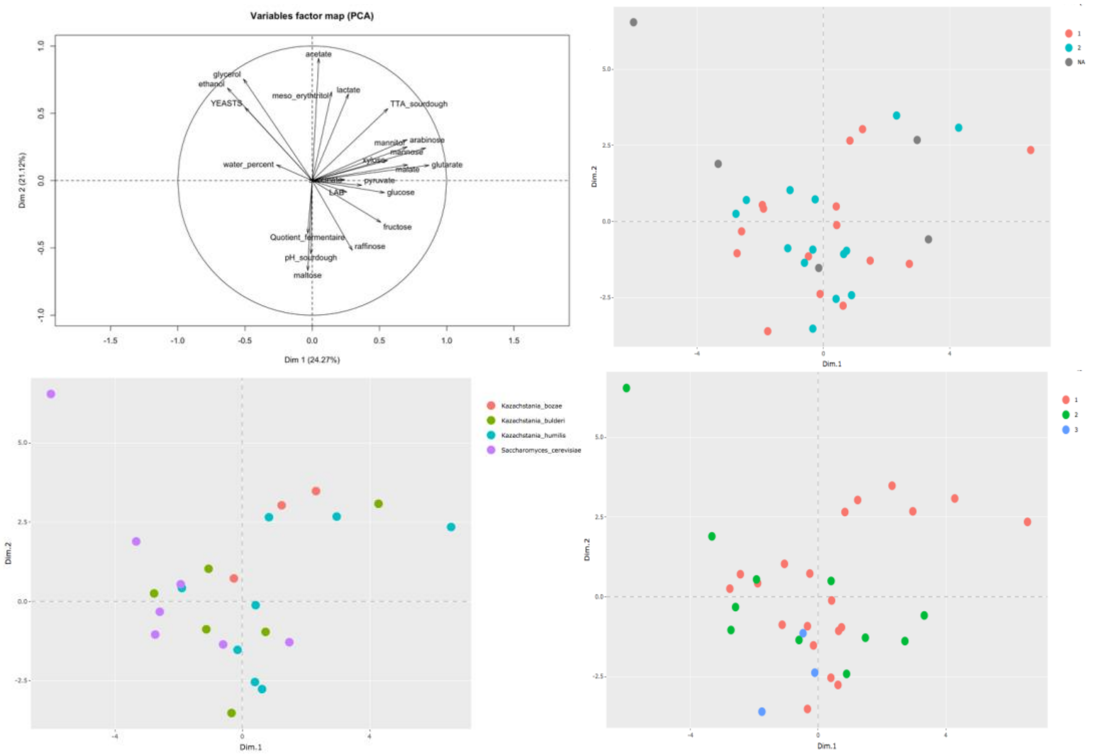


**Fig. S4**. Principal component analysis of 34 sourdoughs based on the quantitative variation of yeast density, bacteria density, sourdough pH, Total Titrable Acidity (TTA), sourdough concentrations of seven sugars (maltose, glucose, fructose, raffinose, arabinose, mannose, xylose), four alcohols (glycerol, ethanol, mannitol, meso_erythtritol), and six acids (lactate, acetate, glutarate, pyruvate, malate, succinate) and the fermentative quotient (lactate over acetate ratio). The correlation between variables are presented on the left while the other figure shows the projection of sourdoughs on the first two axes representing 48% of the variation. Depending on the figures, the sourdoughs are coloured according to the bread-making practice group of the baker (top right), the fungal dominant species (bottom left) or the PcoA microbial community group (bottom right).

**Table S1.** Description of bakers. Code of each baker, location (French department. Region), organic process (yes/no), collection date of sourdoughs and references.

| Bakers (code) | French departement | Region | Organic | Collection_date | References of other published articles with these bakers |
|---|---|---|---|---|---|
| B01 | Vienne | Nouvelle-Aquitaine | Yes | Nov. 2013 | Lhomme et al., 2016; Urien et al., 2019 |
| B02 | Lot-et-Garonne | Nouvelle-Aquitaine | Yes | Dec. 2013 | Lhomme et al., 2016; Urien et al., 2019 |
| B03 | Isère | Auvergne-Rhône-Alpes | Yes | Dec. 2013 | Lhomme et al., 2016; Urien et al., 2019 |
| B04 | Loire-Atlantique | Pays-de-la-Loire | Yes | Jan. 2013 | Lhomme et al., 2016; Urien et al., 2019 |
| B05 | Oise | Hauts-de-France | Yes | Jan. 2013 | Lhomme et al., 2016; Urien et al., 2019 |
| B06 | Aube | Grand-Est | Yes | Mar. 2013 | Lhomme et al., 2016; Urien et al., 2019 |
| B07 | Finistère | Bretagne | Yes | Mar. 2013 | Lhomme et al., 2016; Urien et al., 2019 |
| B08 | Loire-Atlantique | Pays-de-la-Loire | Yes | Mar. 2013 | Lhomme et al., 2016; Urien et al., 2019 |
| B09 | Essonne | Ile-de-France | Yes | Apr. 2013 | Lhomme et al., 2016; Urien et al., 2019 |
| B10 | Alpes-de-Haute-Provence | Provence-Alpes-Côte-d'Azur | Yes | Apr. 2013 | Lhomme et al., 2016; Urien et al., 2019 |
| B11 | Saône-et-Loire | Bourgogne-Franche-Comte | Yes | Apr. 2013 | Lhomme et al., 2016; Urien et al., 2019 |
| B12 | Ille-et-Vilaine | Bretagne | Yes | Apr. 2013 | Lhomme et al., 2016; Urien et al., 2019 |
| B13 | Doubs | Bourgogne-Franche-Comte | Yes | Apr. 2013 | Lhomme et al., 2016; Urien et al., 2019 |
| B14 | Haut-Rhin | Grand-Est | Yes | Apr. 2013 | Lhomme et al., 2016; Urien et al., 2019 |
| B15 | Hérault | Occitanie | Yes | Jan. 2015 | Michel et al., 2016 |
| B16 | Yvelines | Ile-de-France | Yes | Jan. 2015 | Michel et al., 2016 |
| B17 | Somme | Hauts-de-France | Yes | Jan. 2015 | Michel et al., 2016 |
| B18 | Paris | Ile-de-France | Yes | Jan. 2015 | Michel et al., 2016 |
| B19 | Aveyron | Occitanie | Yes | Feb. 2015 | Michel et al., 2016 |
| B20 | Alpes-Maritimes | Provence-Alpes-Côte-d'Azur | Yes | Feb. 2015 | Michel et al., 2016 |
| B21 | Morbihan | Bretagne | Yes | Feb. 2015 | Michel et al., 2016 |
| B22 | Lot-et-Garonne | Nouvelle-Aquitaine | Yes | Feb. 2015 | Michel et al., 2016 |
| B23 | Bas-Rhin | Grand-Est | Yes | Apr. 2015 | Michel et al., 2016 |
| B24 | Vaucluse | Provence-Alpes-Côte-d'Azur | Yes | Feb. 2015 | Michel et al., 2016 |
| B25 | Haut-Rhin | Grand-Est | Yes | Mar. 2015 | Michel et al., 2016 |
| B26 | Côtes-d'Armor | Bretagne | Yes | Mar. 2015 | Michel et al., 2016 |
| B27 | Puy-de-Dôme | Auvergne-Rhône-Alpes | Yes | Mar. 2015 | Michel et al., 2016 |
| B28 | Paris | Ile-de-France | Yes | Mar. 2015 | Michel et al., 2016 |
| B29 | Aude | Occitanie | Yes | Apr. 2015 | Michel et al., 2016 |
| B30 | Savoie | Auvergne-Rhône-Alpes | Yes | Apr. 2015 | Michel et al., 2016 |
| B31 | Haut-Rhin | Grand-Est | Yes | Apr. 2015 | |
| B32 | Saône-et-Loire | Bourgogne-Franche-Comte | Yes | Mar. 2015 | |
| B33 | Ille et Vilaine | Bretagne | Yes | Feb. 2016 | |
| B34 | Hérault | Occitanie | Yes | Oct. 2017 | |
| B40 | Paris | Ile-de-France | No | May 2015 | |
| B41 | Lozère | Occitanie | No | Jun. 2015 | |
| B42 | Loire | Auvergne-Rhône-Alpes | No | Jul. 2015 | |
| B43 | Gard | Occitanie | No | Jul. 2015 | |
| B44 | Loire-Atlantique | Pays-de-la-Loire | No | Mar. 2016 | |

**Table S2.** Oligonucleotides primers used for PCR amplification and sequencing

| Region amplification | Primers | Sequence | Reference |
|---|---|---|---|
| *ITS (ITS1- 5.8S - ITS2)* | ITS1F | 5’-CTTGGTCATTTAGAGGAAGTAA-3’ | White *et al.*, 1990 |
| | ITS4 | 5’-TCCTCCGCTTATTGATATGC-3’ | |
| | NSA3 | 5’- AAACTCTGTCGTGCTGGGGATA-3’ | Martin et Rygiewicz. 2005 |
| | NLB4 | 5’- GGATTCTCACCCTCTATGAC-3’ | |
| *ITS1* | NSA3 | 5’- AAACTCTGTCGTGCTGGGGATA-3’ | Martin et Rygiewicz. 2005 |
| | 58A2R | 5’- CTGCGTTCTTCATCGAT -3’ | |
| | ITS1F | 5’-CTTGGTCATTTAGAGGAAGTAA-3’ | White *et al.*, 1990 |
| | ITS2 | 5'-GCTGCGTTCTTCATCGATGC-3' | Buée *et al.*, 2009 |
| LSU (D1D2) | NL1 | 5’-GCATATCAATAAGCGGAGGAAAAG-3’ | O’Donnell *et al.*, 1993 |
| | NL4 | 5’-GGTCCGTGTTTCAAGACGG-3’ | |
| *RPB1* | RPB1-Af | 5’-GARTGYCCDGGDCAYTTYGG-3’ | Matheny *et al.*, 2002 |
| | RPB1-Cr | 5’-CCNGCDATNTCRTTRTCCATRTA-3’ | |
| *RPB2* | RPB2-5F | 5’-GAYGAYMGWGATCAYTTYGG-3’ | Liu *et al.*, 1999 |
| | RPB2-7cR | 5’-CCCATRGCTTGYTTRCCCAT-3’ | |
| LSU | LSU-LROR | 5’- ACCCGCTGAACTTAAGC-3’ | Vilgalys and Hester. 1990 |
| | LSU-LR5 | 5’- TCCTGAGGGAAACTTCG-3’ | |
| *Act1* | CA5R | 5’-GTGAACAATGGATGGACCAGATTCGTCG-3’ | Kan *et al.*, 1993 |
| | CA14 | 5’-AACTGGGATGACATGGAGAAGATCTGGC-3’ | |
| TEF | YTEF-1G | 5’-GGTAAGGGTTCTTTCAAGTACGCTTGGG-3’ | Kurtzman and Robnett. 2003 |
| | YTEF-6G | 5’-CGTTCTTGGAGTCACCACAGACGTTACCTC-3’ | |
| *GDH1* | GDH1-F | 5’- TGGAAATGAGCGGAAGAAGAAAGC-3’ | Peris *et al.*, 2014 |
| | GDH1-R | 5’- CTGTAGGCACCGAACAAGTAACC-3’ | Nguyen and Gaillardin. 2005 |
| *FSY1* | FSY1-F | 5’- GGATCYTCRACAAGCGTTTCTC-3’ | Libkind. Hittinger *et al.*, 2011 |
| | FSY1-R | 5’- AAGGCAAACAYGTAAAGCAAAG-3’ | |
| *URA3* | URA3-F | 5’- GCGCCCCTTTCTCTTATGTT-3’ | Peris *et al.*, 2014 |
| | URA3-R | 5’- CATTTGCTTTTGTTCCACCA-3’ | |
| *DRC1* | DRC1-F | 5’- GCAATTAAGGGAGGCAATAAAAGTC-3’ | Peris *et al.*, 2014 |
| | DRC1-R | 5’- CAACACAAGGGCACTCATAATC-3’ | |
| *MET2* | MET2-F | 5’- CGAAAACGCTCCAAGAGCTGG-3’ | Sampaio and Gonçalves. 2008 |
| | MET2-R | 5’- GACCACGATATGCACCAGGCAG-3’ | |

**Table S3.** Alpha diversity indexes. Number of observed species, Chao1 index, effective number of species from Shannon and Simpson indexes. True Shannon diversity indexes were calculated with $exp^{Shannon\ index}$, and True Simpson diversity indexes with $\frac{1}{1-Simpson\ index}$.

| | | | Effective number of species from | |
|---|---|---|---|---|
| | Observed | Chao1 | Shannon | Simpson |
| Sourdough_B01 | 5 | 6 | 1.03 | 1.01 |
| Sourdough_B10 | 1 | 1 | 1 | 1 |
| Sourdough_B11 | 2 | 2 | 1.0 | 1 |
| Sourdough_B12 | 10 | 13 | 1.16 | 1.06 |
| Sourdough_B13 | 6 | 8 | 1.06 | 1.02 |
| Sourdough_B14 | 25 | 26 | 3.42 | 1.85 |
| Sourdough_B15 | 3 | 4 | 1.02 | 1 |
| Sourdough_B16 | 15 | 15 | 2.52 | 1.92 |
| Sourdough_B17 | 11 | 14 | 1.18 | 1.06 |
| Sourdough_B18 | 20 | 20 | 4.39 | 2.67 |
| Sourdough_B19 | 14 | 17 | 1.14 | 1.04 |
| Sourdough_B02 | 16 | 24 | 1.53 | 1.20 |
| Sourdough_B20 | 20 | 21 | 3.83 | 2.72 |
| Sourdough_B21 | 12 | 13 | 1.19 | 1.06 |
| Sourdough_B22 | 4 | 4 | 1.04 | 1.01 |
| Sourdough_B23 | 9 | 11 | 1.06 | 1.02 |
| Sourdough_B24 | 13 | 16 | 1.14 | 1.04 |
| Sourdough_B25 | 21 | 21 | 3.44 | 1.86 |
| Sourdough_B26 | 6 | 7 | 1.02 | 1.01 |
| Sourdough_B27 | 16 | 16 | 1.83 | 1.29 |
| Sourdough_B28 | 14 | 19 | 1.43 | 1.16 |
| Sourdough_B29 | 17 | 17 | 2.33 | 1.59 |
| Sourdough_B03 | 10 | 15 | 1.10 | 1.03 |
| Sourdough_B30 | 18 | 18 | 2.30 | 1.43 |
| Sourdough_B31 | 5 | 5 | 1.33 | 1.16 |
| Sourdough_B32 | 20 | 21 | 1.57 | 1.18 |
| Sourdough_B33 | 16 | 19 | 1.53 | 1.2 |
| Sourdough_B04 | 18 | 21 | 1.4 | 1.16 |
| Sourdough_B40 | 20 | 22 | 2.25 | 1.54 |
| Sourdough_B41 | 8 | 8 | 2.11 | 2.03 |
| Sourdough_B42 | 26 | 40 | 6.79 | 5 |
| Sourdough_B43 | 26 | 29 | 2.06 | 1.42 |
| Sourdough_B44 | 25 | 32 | 4.11 | 2.95 |

| | | | | |
|---|---|---|---|---|
| Sourdough_B05 | 9 | 30 | 1.09 | 1.03 |
| Sourdough_B06 | 6 | 9 | 1.02 | 1.01 |
| Sourdough_B07 | 14 | 24 | 1.76 | 1.42 |
| Sourdough_B08 | 4 | 4 | 1.06 | 1.02 |
| Sourdough_B09 | 22 | 25 | 2.87 | 1.99 |

**Table S4.** Univariate permutational Manova analysis performed on the Unifrac distances between the sourdoughs of 30 bakers which had less than 8 missing values among the 29 practices variables. Bakers 6, 10, 19 et 20 were excluded as they had more than 8 missing values.

| var | Df | SumsOfSqs | MeanSqs | F.Model | R2 | Pr(>F) | nb | p.adj |
|---|---|---|---|---|---|---|---|---|
| Type baker | 1 | 0.11 | 0.11 | 1.28 | 0.04 | 0.23 | 30 | 0.46 |
| Milling | 1 | 0.18 | 0.18 | 2.16 | 0.07 | 0.11 | 30 | 0.29 |
| Bran | 1 | 0.05 | 0.05 | 0.59 | 0.02 | 0.53 | 30 | 0.62 |
| Conservation temperature sourdough | 3 | 0.32 | 0.11 | 1.23 | 0.12 | 0.25 | 30 | 0.46 |
| Chief origin | 2 | 0.53 | 0.26 | 3.54 | 0.21 | 0.01 | 30 | 0.09 |
| Water origin | 4 | 0.51 | 0.13 | 1.55 | 0.20 | 0.13 | 30 | 0.31 |
| Flour origin | 2 | 0.15 | 0.08 | 0.86 | 0.06 | 0.47 | 30 | 0.62 |
| Yeast use bakery | 1 | 0.47 | 0.47 | 6.43 | 0.19 | 0.00 | 30 | 0.09 |
| Kneading method | 2 | 0.39 | 0.20 | 2.45 | 0.15 | 0.04 | 30 | 0.18 |
| Mill type | 3 | 0.49 | 0.16 | 2.06 | 0.19 | 0.06 | 30 | 0.22 |
| Variety cereals dough | 2 | 0.41 | 0.21 | 2.62 | 0.16 | 0.05 | 30 | 0.18 |
| Kg bread week | 3 | 0.65 | 0.22 | 3.00 | 0.26 | 0.02 | 30 | 0.11 |
| Age sourdough collection | 2 | 0.16 | 0.08 | 0.91 | 0.07 | 0.45 | 29 | 0.62 |
| Age sourdough year | 3 | 0.69 | 0.23 | 3.49 | 0.30 | 0.01 | 29 | 0.09 |
| Hydratation backslopping | 2 | 0.12 | 0.06 | 0.65 | 0.05 | 0.62 | 28 | 0.70 |
| Backslopping temperature | 2 | 0.23 | 0.11 | 1.31 | 0.09 | 0.27 | 30 | 0.48 |
| Backslopping period | 2 | 0.07 | 0.04 | 0.43 | 0.03 | 0.81 | 28 | 0.81 |
| Temperature dough end kneading | 2 | 0.13 | 0.07 | 0.77 | 0.06 | 0.53 | 26 | 0.62 |
| Sourdough weight dough | 2 | 0.10 | 0.05 | 0.54 | 0.04 | 0.75 | 30 | 0.78 |
| Flour weight dough | 1 | 0.13 | 0.13 | 1.48 | 0.05 | 0.20 | 30 | 0.43 |
| Water weight dough | 2 | 0.34 | 0.17 | 2.07 | 0.13 | 0.08 | 30 | 0.25 |
| Salt weight dough | 1 | 0.04 | 0.04 | 0.40 | 0.01 | 0.75 | 29 | 0.78 |
| Total fermentation period | 2 | 0.40 | 0.20 | 2.54 | 0.16 | 0.04 | 30 | 0.18 |
| First fermentation | 1 | 0.06 | 0.06 | 0.72 | 0.03 | 0.46 | 29 | 0.62 |
| Final fermentation | 1 | 0.18 | 0.18 | 2.06 | 0.07 | 0.11 | 29 | 0.29 |
| Nb backslopping week | 1 | 0.09 | 0.09 | 1.02 | 0.04 | 0.39 | 30 | 0.60 |
| Nb backslopping before use | 2 | 0.18 | 0.09 | 1.02 | 0.07 | 0.35 | 30 | 0.57 |
| Nb breadmaking week | 2 | 0.15 | 0.07 | 0.83 | 0.06 | 0.50 | 30 | 0.62 |

**Table S5.** Summary statistics on each of the quantitative variable calculated over all sourdoughs. Yeasts and Lactic Acid Bacteria (LAB) density are expressed in cells per gram of sourdough. Total Titrable Acid (TTA) in mL of NaOH 0.1M added to reach pH8.3. Water content is expressed in percent. Sugar, alcohol, organic acid concentrations are expressed in mg / gr of sourdough.

| Quantitative variable | Mean | SD | Median | Min | Max |
|---|---|---|---|---|---|
| YEASTS | 2.87E+07 | 5.76E+07 | 2.00E+07 | 8.09E+04 | 5.81E+08 |
| LAB | 1.17E+09 | 1.16E+09 | 9.35E+08 | 1.44E+07 | 6.80E+09 |
| pH_sourdough | 3.87 | 0.20 | 3.84 | 3.41 | 4.81 |
| TTA_sourdough | 15.93 | 5.36 | 15.60 | 4.52 | 31.30 |
| water (%) | 52.34 | 9.43 | 49.87 | 39.48 | 91.03 |
| Maltose | 26.49 | 15.25 | 23.30 | 1.08 | 57.93 |
| Glucose | 24.97 | 9.29 | 26.03 | 6.09 | 41.72 |
| Ethanol | 7.54 | 5.01 | 6.76 | 1.08 | 27.05 |
| Lactate | 7.23 | 1.85 | 7.27 | 1.94 | 11.43 |
| Mannitol | 5.09 | 2.52 | 5.07 | 0.00 | 12.36 |
| Raffinose | 3.34 | 2.56 | 2.97 | 0.00 | 9.82 |
| Fructose | 2.52 | 1.87 | 2.04 | 0.32 | 10.31 |
| Glycerol | 2.17 | 1.30 | 1.96 | 0.27 | 6.40 |
| Acetate | 1.65 | 0.64 | 1.63 | 0.70 | 3.83 |
| Arabinose | 0.93 | 0.60 | 0.79 | 0.17 | 3.76 |
| Mannose | 0.92 | 0.34 | 0.86 | 0.51 | 2.61 |
| Xylose | 0.69 | 0.36 | 0.63 | 0.00 | 1.93 |
| Succinate | 0.62 | 0.22 | 0.56 | 0.27 | 1.26 |
| Meso_erythtritol | 0.28 | 0.08 | 0.26 | 0.12 | 0.50 |
| Malate | 0.27 | 0.12 | 0.24 | 0.02 | 0.65 |
| Glutarate | 0.04 | 0.02 | 0.04 | 0.01 | 0.09 |
| Pyruvate | 0.03 | 0.01 | 0.02 | 0.01 | 0.07 |
| Fermentative quotient | 3.15 | 1.03 | 2.97 | 1.37 | 6.33 |

**Table S6.** Confident intervals of the ratio between sourdough and non-sourdough strains for *K. bulderi* (B) and *K. humilis* (H) and each quantitative variable (t1g, tVmax, Vmax, CO2mx, Cellt27, Mortality).

| estimate | lwr | upr | trait | Species | p.value | p.adj |
|---|---|---|---|---|---|---|
| **1.07471296434514** | 0.945568940277162 | 1.22149523586612 | tVmax | B | 0.269979267696504 | 0.539958535393007 |
| **0.767703695980246** | 0.665742272844319 | 0.885280969621636 | tVmax | H | 0.000277018785404315 | 0.00166211271242589 |
| **0.92725089908944** | 0.794574508107426 | 1.08208131659055 | Vmax | B | 0.337715651630287 | 0.578941117080492 |
| **1.21600473680224** | 1.02451724325925 | 1.44328221867839 | Vmax | H | 0.0252850365667922 | 0.0758551097003765 |
| **0.946325862856667** | 0.9022962364222 | 0.992504016488417 | CO2max | B | 0.0232383975488227 | 0.0758551097003765 |
| **1.00747139218045** | 0.95838424745388 | 1.05907271405863 | CO2max | H | 0.770228536886918 | 0.781824893399713 |
| **1.05847085933614** | 0.826618920819044 | 1.35535315227686 | Cellt27 | B | 0.652363541323959 | 0.781824893399713 |
| **0.963211563861061** | 0.738783632601456 | 1.2558162847879 | Cellt27 | H | 0.781824893399713 | 0.781824893399713 |
| **0.883655742699433** | 0.634194385473931 | 1.23124311644948 | Mortality | B | 0.464886001170052 | 0.636538523489544 |
| **1.29820938325557** | 0.911708780173879 | 1.84855914456738 | Mortality | H | 0.147799138902406 | 0.354717933365775 |
| **1.04166853707392** | 0.930722615588945 | 1.16583966367155 | t1g | B | 0.477403892617158 | 0.636538523489544 |
| **0.775149043181731** | 0.684374479226676 | 0.877963830306038 | t1g | H | 6.12225373166631e-05 | 0.000734670447799957 |

**Additional data set and scripts**

Michel Elisa, Masson Estelle, Bubendorff Sandrine, … Sicard Delphine. (2022). Artisanal and farmers bread-making practices differently shape fungal species community diversity in French sourdoughs (Version 2) [Data set]. Zenodo. http://doi.org/10.5281/zenodo.5849058
